# Top-down control of exogenous attentional selection is mediated by beta coherence in prefrontal cortex

**DOI:** 10.1101/2023.01.11.523664

**Authors:** Agrita Dubey, David A. Markowitz, Bijan Pesaran

## Abstract

Salience-driven exogenous and goal-driven endogenous attentional selection are two distinct forms of attention that guide selection of task-irrelevant and task-relevant targets in primates. During conflict i.e, when salience and goal each favor the selection of different targets, endogenous selection of the task-relevant target relies on top-down control. Top-down attentional control mechanisms enable selection of the task-relevant target by limiting the influence of sensory information. Although the lateral prefrontal cortex (LPFC) is known to mediate top-down control, the neuronal mechanisms of top-down control of attentional selection are poorly understood. Here, using a two-target free-choice luminance-reward selection task, we demonstrate that visual-movement neurons and not visual neurons or movement neurons encode exogenous and endogenous selection. We then show that coherent-beta activity selectively modulates mechanisms of exogenous selection specifically during conflict and consequently may support top-down control. These results reveal the VM-neuron-specific network mechanisms of attentional selection and suggest a functional role for beta-frequency coherent neural dynamics in the modulation of sensory communication channels for the top-down control of attentional selection.

## Introduction

In primates, selection of task-relevant targets is guided by goal-driven (“top-down”) endogenous attentional processes whereas selection of task-irrelevant distractors is guided by salience-driven exogenous (“bottom-up”) attentional processes (Awh et al., 2006; Carrasco et al., 2004; Corbetta and Shulman, 2002; Corbetta et al., 1998; Moore and Zirnsak, 2017; Theeuwes, 2010). Exogenous selection is fast and occurs earlier in time whereas endogenous selection is slow and occurs later in time (Buschman and Miller, 2007; Dugué et al., 2020; Markowitz et al., 2011; Theeuwes, 2010). Therefore, trial-by-trial flexible selection behavior depends on the dynamic interplay between exogenous and endogenous attentional mechanisms (Buschman and Miller, 2007; Markowitz et al., 2011; Pesaran et al., 2021; Theeuwes, 2010). However, how attentional selection is controlled when exogenous and endogenous attentional mechanisms are in conflict remains unclear. How is the task-relevant target selected when in conflict with salient target?

Endogenous attentional selection relies on a top-down control process that enables the selection of task-relevant targets by limiting the influences of automatic-salience selection (Anderson and Weaver, 2009; Miller and Cohen, 2001; Womelsdorf and Everling, 2015). Neural mechanisms that support top-down control are distributed throughout the fronto-parietal regions and rely heavily on the lateral prefrontal cortex (LPFC) (Buschman and Kastner, 2015; Buschman and Miller, 2007; Miller, 2000; Miller and Cohen, 2001; Moore and Armstrong, 2003; Paneri and Gregoriou, 2017; Suzuki and Gottlieb, 2013) as evident from lesion experiments (Buckley et al., 2009; Gregoriou et al., 2014; Petrides, 2005; Rossi et al., 2007; Rushworth et al., 2005). Thus, LPFC-mediated top-down control mechanisms may support selection of the task-relevant target when in conflict with task-irrelevant salient target.

Information flow about task-relevant and irrelevant targets during conflict must be mediated by multiregional communication and, specifically, competition between convergent information streams. Since exogenous attentional selection is fast and processes sensory streams of information while endogenous attentional selection is slow and processes information about goals, each attentional process operates across distinct neural pathways, i.e. communication channels (**Fig 1A**). Consequently, selective filtering of information flow across sensory and reward-based communication channels may support the top-down control of attentional selection.

**Figure 1:**
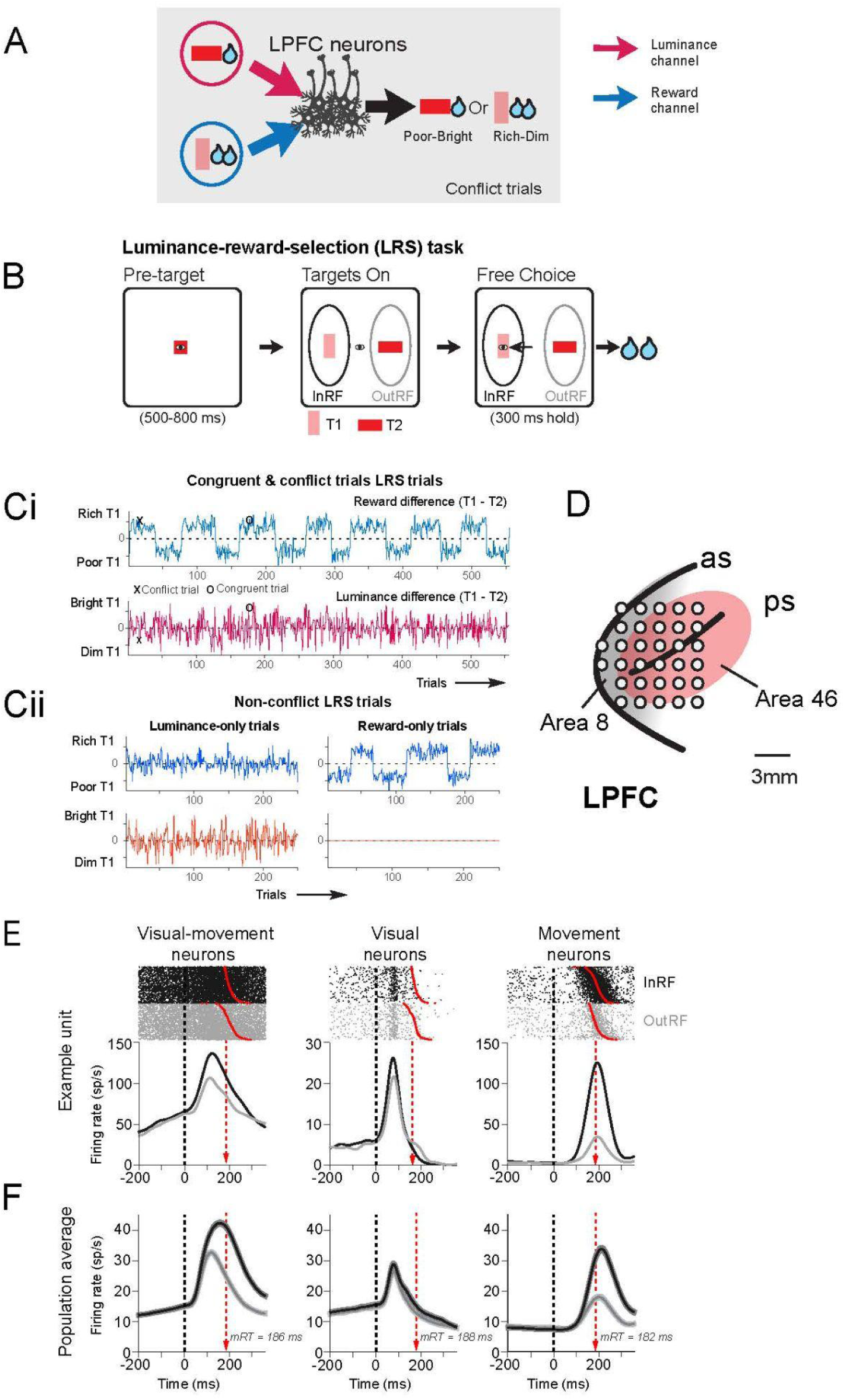
Experimental design to investigate the mechanisms of attentional selection. **(A)**The mechanisms of attentional selection involve filtering information flow across communication channels. The luminance channel communicates the task-irrelevant sensory stream of information while the reward channel communicates task-relevant information about the goal to LPFC. During conflict, the luminance and reward communication channels compete to guide exogenous or endogenous selection. The top-down control of attentional selection may guide exogenous or endogenous selection by filtering either or both of the luminance and reward channels. **(B)** Luminance-reward-selection (LRS) task. At the start of each LRS trial, each monkey maintained fixation on a visual fixation target presented at the center of the screen for a 500-800 ms baseline period. After the baseline, the fixation target was extinguished and two peripheral saccade targets, a horizontal bar and a vertical bar, were presented at random relative locations on a 10 deg circle in the visual periphery, with a minimum angular separation of 90 degrees. Each monkey made a saccade to one of the visual targets immediately following target presentation to earn a water reward. **(Ci)** congruent and conflict LRS trials - Mean value of reward associated with each target is varied in blocks of 40-70 trials (top panel). Luminance value associated with each target is randomly selected on each trial (bottom panel). **(Cii)** Same as Ci except for luminance-only (left panel) and reward-only (right panel) non-conflict LRS trials. **(D)** Neural recording locations over lateral prefrontal cortex. White dots indicate electrode penetration sites. Area 8 (grey) and 46 (red) indicated based on Petrides and Pandya (Petrides and Pandya, 1994). Sulcal landmarks: *as*, arcuate sulcus; *ps*, principal sulcus.**(E)** Spike rasters and peri-stimulus time histograms (PSTHs) of an example visual-movement (VM), visual and movement neuron for congruent and conflict LRS trials aligned to target presentation. Dark-grey traces denote selection into the RF; light-grey traces denote selection out of the RF. Red dots denote saccade reaction time of each trial. Dotted lines denote target onset. **(F)** Population average firing rates of VM (N=139, left), visual (N=57, middle) and movement (N=65, right) neurons on congruent and conflict LRS trials when target selection was InRF (dark-grey) and OutRF (light-grey). The s.e.m of firing rates is shaded in lighter shades. Red arrows denote average reaction times.

Neuronal coherence measured by local field potential (LFP) activity in specific frequency bands reveals the correlations in the timing of neural activity across populations of neurons (Pesaran et al., 2018), and is generally interpreted in terms of multiregional communication (Hagan and Pesaran, 2022; Staudigl et al., 2022; Voloh and Womelsdorf, 2016). Many studies highlight the importance of multiregional communication and neuronal coherence to attentional selection. Attentional selection involves interactions between populations of LPFC neurons (Fiebelkorn and Kastner, 2021; Panichello and Buschman, 2021). In LPFC, cue-triggered LFP activity in the beta-frequency (15-35 Hz) band reflects exogenous selection (Buschman and Miller, 2007; Fiebelkorn and Kastner, 2021). Beta frequency activity after the cue also reflects endogenous selection and suppression of sensory information during working memory and attention tasks (Antzoulatos and Miller, 2016; Buschman et al., 2012; Hanslmayr et al., 2014; Lundqvist et al., 2018; Miller et al., 2018; Schmidt et al., 2019; Spitzer and Haegens, 2017). This suggests that beta frequency neuronal coherence may support top-down control and the trial-by-trial interplay between endogenous and exogenous attentional selection during conflict. However, prior work has not dissociated endogenous and exogenous selection during conflict to understand how beta frequency coherence biases information flow across communication channels to guide attentional selection. Whether beta frequency neural coherence acts on communication channels carrying salience-driven or goal-directed information is not known.

Here, we test the neural mechanisms of endogenous attentional selection by recording local-field potential (LFP) and spiking activity from neurons across LPFC of macaque monkeys. We study top-down control of attentional selection using a simple two-target, free-reaction time, luminance-reward-selection (LRS) task. The LRS task independently manipulated relative luminance and reward values of the two targets to yield a subset of conflict trials. On these trials, sensory and reward drives favored different targets so that exogenous and endogenous attentional selection processes were in conflict. We therefore compared neural activity and behavior between conflict and non-conflict trials to better understand the top-down control of attentional selection and the role of beta-frequency neuronal coherence.

## Results

We trained two Rhesus macaque monkeys (*macaca mulatta*) to perform a **luminance-reward-selection (LRS) task** (**Fig 1B**, see **Methods**). The LRS task dissociates exogenous and endogenous attentional selection by independently manipulating reward and luminance (**Fig 1Ci**). Trial-by-trial independent manipulation of luminance and reward values yielded either congruent or conflict set of trials. On congruent trials, luminance and reward value was high for one target- **Rich-Bright** and low for other target **Poor-Dim**. On conflict trials, luminance and reward drives were in conflict, one target was **Rich-Dim** while the other was **Poor-Bright (Fig 1Ci)**. On a subset of trials, the LRS task featured non-conflict luminance-only trials and reward-only trials. On luminance-only trials, the relative reward values associated with two targets were kept similar across blocks. On reward-only trials, the luminance values of two targets were kept the same for blocks **(Fig 1Cii)**. Monkeys were required to make a saccade to one of the targets immediately following target presentation, to earn a water reward.

We recorded local field potential (LFP) and single unit spiking activity from 32 electrodes in LPFC during LRS task performance (**Fig 1D**. Monkey 1: N=39 sessions, Monkey 2: N=42 sessions) yielding 409 task-responsive single units (M1: 179; M2: 230 neurons, **Methods**). We also recorded the activity of each neuron during the performance of an oculomotor delayed response (ODR) task involving a single visual target (ODR trials, see **Methods**). Due to the obligatory relationship between oculomotor behavior and visual-spatial attention (Kowler et al., 1995), requiring a saccadic eye movement revealed the spatial locus of visual attention on each LRS trial. Testing responses during the ODR task revealed neurons with delay representations that could guide attentional selection.

Consistent with earlier work, LPFC neuron activity increased in a spatially-selective manner during the ODR trials (Funahashi et al., 1989). (see **Methods**). Of 409 neurons, 261 neurons were ODR-task-responsive with an excitatory response field (64%). The majority of excitatory ODR-task-responsive LPFC neurons showed elevated firing activity in response to both target onset and the saccadic response, which we term visual-movement (VM) neurons (N=139, 53%; **Supp Fig 1)**. A substantial minority of excitatory response field neurons displayed increased firing in response to target onset alone, termed visual neurons (N=57, 22%; **Supp Fig 2**), or around the saccadic eye movement and not target onset, termed movement neurons (N=65, 25%; **Supp Fig 3**).

**Figure 2:**
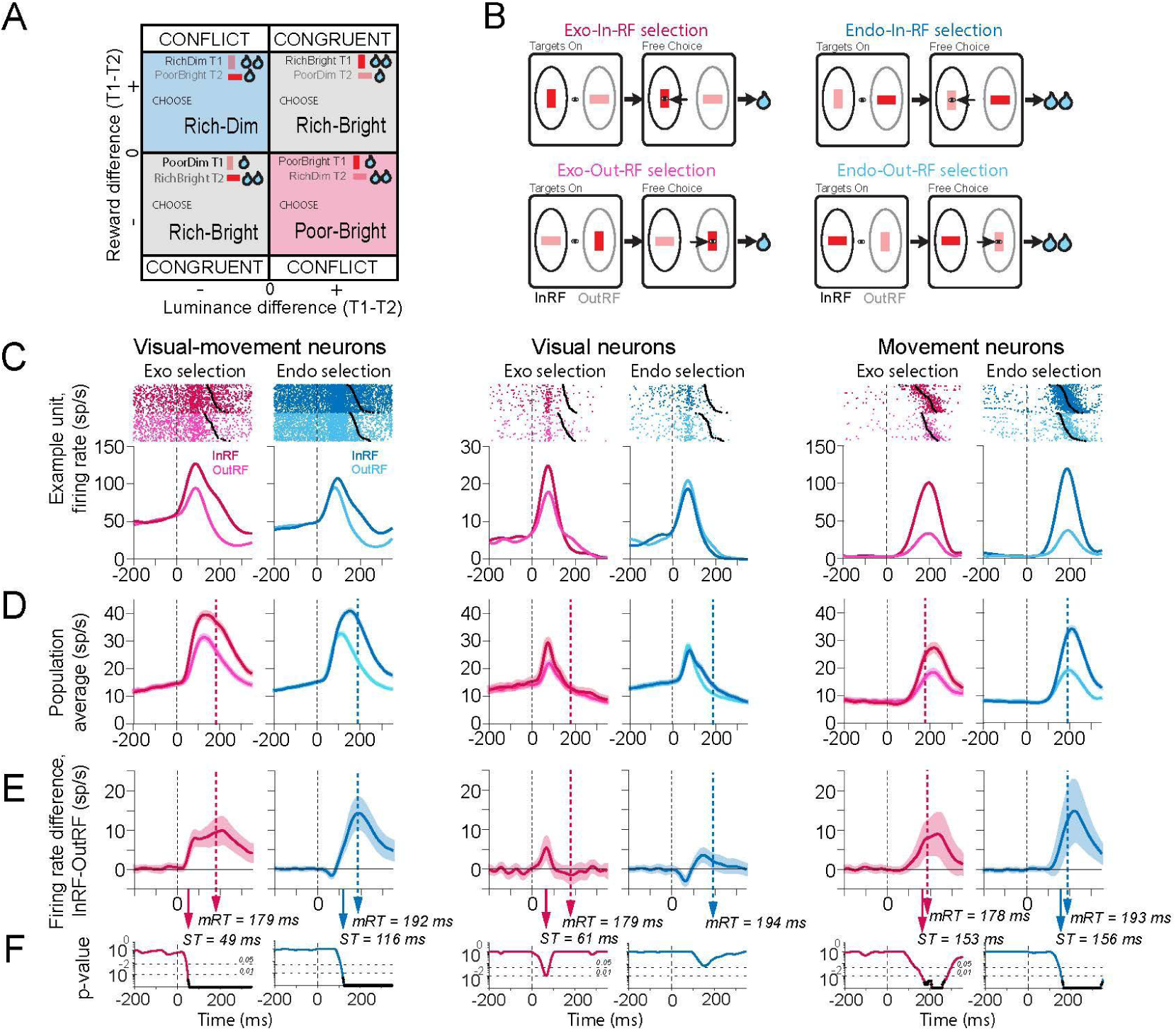
Visual-movement (VM) neurons reflect exogenous and endogenous attentional selection. **(A)**Choice behavior plotted as a function of luminance and reward difference between the two targets. LRS task yields two major sets of trials -congruent and conflict. On congruent trials, luminance and reward value is high for one target-**Rich-Bright** and low for other target **Poor-Dim**. When both luminance and reward drive are in congruent, they both favored the selection of the Rich-Bright target. On conflict trials, one target has high-reward and low-luminance **Rich-Dim** while the other has low-reward and high-luminance **Poor-Bright.** When luminance and reward drive are in conflict, luminance driven choices result in exogenous (**Exo**) selection of the Poor-Bright target and reward driven choices result in endogenous (**Endo**) selection of the Rich-Dim target. Note that target properties Rich/Poor-Bright/Dim are independent of the orientation of the target, i.e, T1 can be Rich-Dim on one trial, Rich-Bright on next trial and Poor-Bright on a given trial of the next block. Same applies for T2. **(B)** Schematic of exogenous and endogenous selection. On **Exo-InRF** trials selected Poor-Bright target is in the RF while on **Exo-OutRF** selected Poor-Bright target is out of the RF. On **Endo-InRF** trials selected Rich-Dim target is in the RF while on **Endo-OutRF** selected Rich-Dim target is out of the RF. **(C)** Spike rasters and PSTHs for exo and endo selection of an example visual-movement, visual and movement neuron on conflict trials shown aligned to the target presentation. Darker traces denote selection into the RF; lighter traces denote selection out of the RF. Black dots denote saccade reaction time of each trial. Dotted lines denote target onset. **(D)** Population average firing rates for VM, visual and movement neurons on Exo-InRF, Exo-OutRF, Endo-InRF and Endo-OutRF conflict trials. The s.e.m of firing rates is shaded in lighter shades. **(E)** Difference in firing rates for selection into and out of the RF for three groups of neurons on conflict trials (top panel). Mean +/- s.e.m. are shown. **(F)** Permutation test p-values against a null hypothesis that there is no difference in InRF and OutRF firing rates are shown in the bottom panel. False-discovery-rate (FDR) corrected p-values for alpha=0.01are shown in black. Arrow represents the selection time (ST) when first time separation becomes significant (VM: Exo ST=49 ms, Endo ST= 116 ms; Visual: Exo ST = 61 ms ; Movement: Exo ST=153 ms, Endo ST=156 ms). Dotted lines denote average reaction time for exogenous (pink) and endogenous (blue) selection.

**Figure 3:**
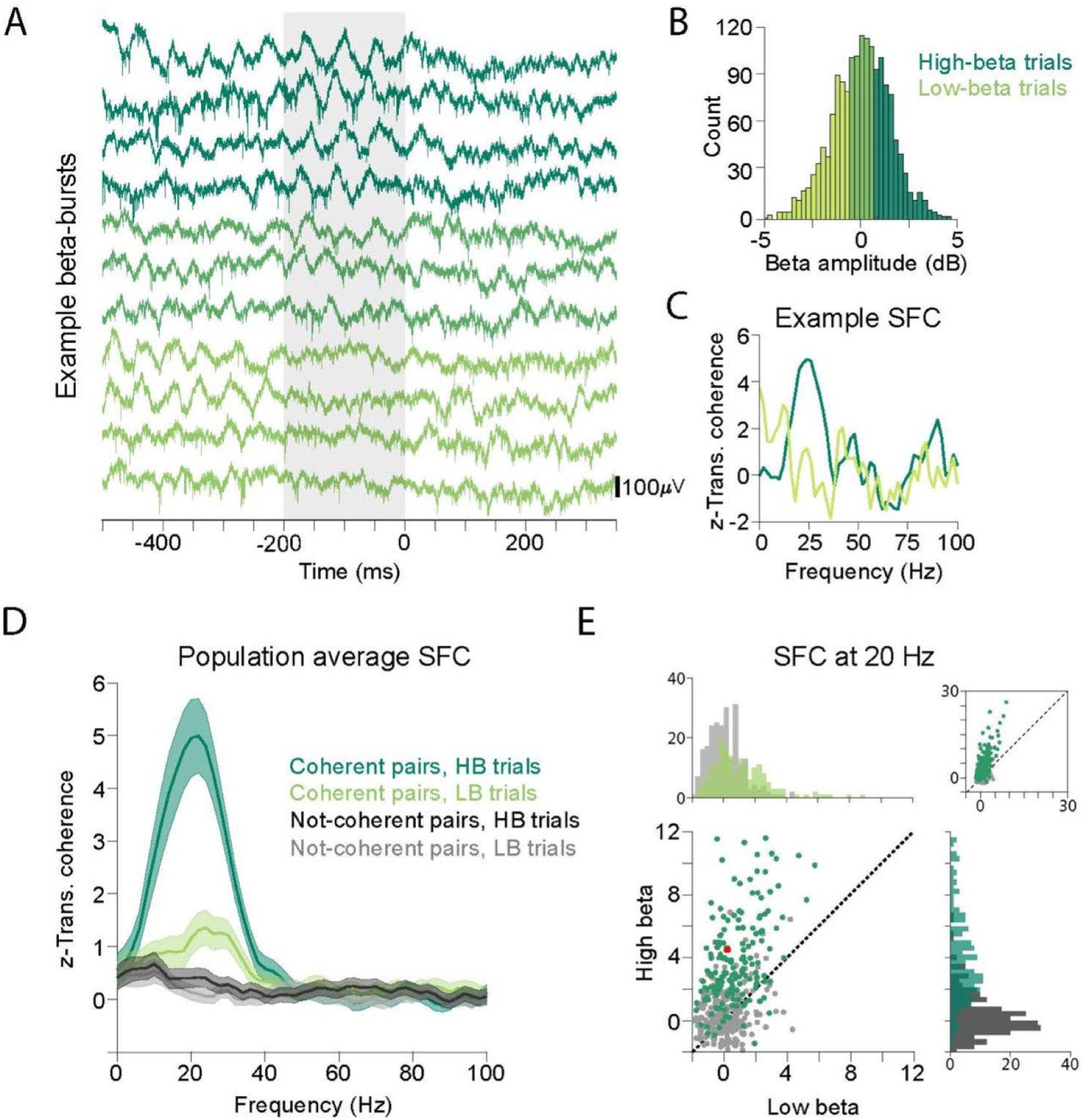
Beta-frequency bursts and coherent neuronal dynamics. **(A)** Raw extracellular recordings at an example recording site from several example trials during LRS task. Shaded area denotes the window of interest used for calculating beta amplitude values. **(B)** Pre-target beta-burst amplitude values across trials at the example site (same as A) on an example experimental session. Dark-green denotes high-beta trials (HB, ∼33% highest beta-bursts); Light-green denotes low-beta trials (LB, ∼33% lowest beta-bursts). **(C)** Spike-field coherence (SFC) between an example unit and field recorded on a neighboring electrode (same as A and B). Darker trace denotes SFC for HB trials while lighter trace denotes SFC for LB trials. **(D)** Population average SFC of coherent pairs (N=176) and not-coherent pairs (N=233). The s.e.m of SFC is shown in lighter shades. **(E)** Scatter plot of HB SFC versus LB SFC at 20 Hz. Plot limits are zoomed in for a better visualization. Inset shows all the SFC electrode pairs. Each dot denotes a recording pair. Red dot denotes the example SFC in C. Marginal histograms denote the SFC distribution for HB and LB trials.

To further analyze the LPFC neuronal population dynamics, we performed a principal component analysis of activity during the luminance-only trials that extracted visual and movement modes. The first two modes represented movement and visual activity respectively and explained ∼75% variability (**Supp. Fig. 4 A,B**). To visualize the LPFC activity on the independent set of LRS task trials in the space of the visual and movement modes for each of the three classes of neurons, we projected visual-movement, visual, and movement firing rates activity to the first two modes (**Supp. Fig. 4C**). While visual-movement neurons showed activity for both visual and movement modes, visual neurons showed activity for the second mode only. Similarly, movement neurons showed activity for the first mode only and not the visual mode. This data-driven analysis supports the classification of dominant neural responses according to the visual and movement modes and LPFC neurons into three groups- VM, visual and movement neurons.

**Figure 4:**
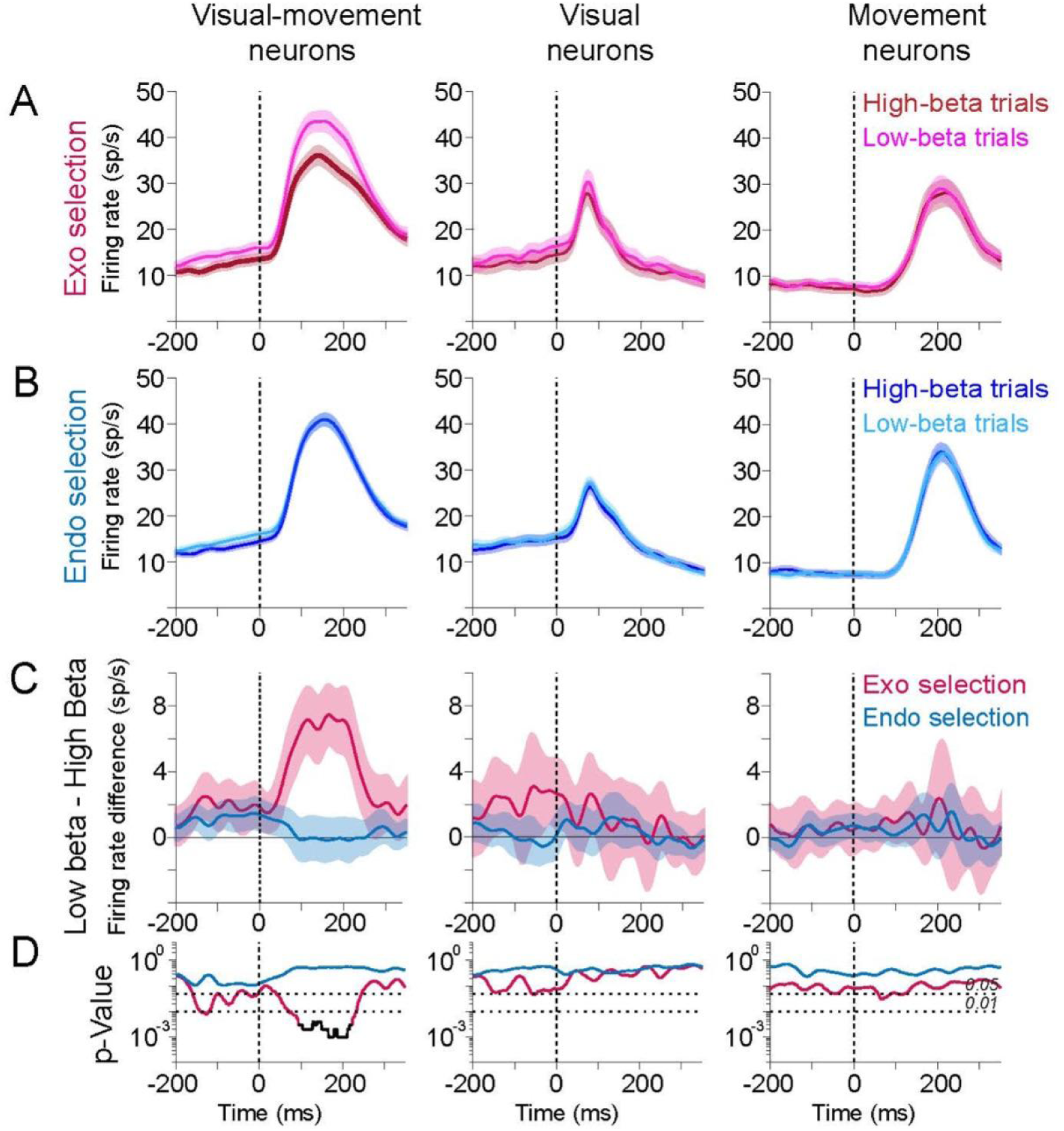
Beta-bursts selectively modulate VM neurons firing activity for exogenous selection during conflict. **(A)** Population average firing rates of the VM, visual and movement neurons for exogenous selection when pre-target beta-burst is high (HB trials, darker traces) and low (LB trials, lighter traces). Mean +/- s.e.m. are shown for InRF conflict trials when selection was in the RF of the units. Dotted lines denote target onset. **(B)** Same as A but for endogenous selection. **(C)** Difference in firing rates for pre-target low and high beta-bursts. Mean +/- s.e.m. are shown for exogenous (red) and endogenous (blue) selection.**(D)** Permutation test p-values under a null hypothesis that there is no difference in firing rates for high-beta and low-beta trials for exogenous selection (red) and endogenous selection (blue). FDR corrected for p-values for alpha=0.01 are shown in black.

We next analyzed responses of these neurons during the LRS task. LPFC neurons involved in attentional selection should fire more spikes on trials when the target in the RF is chosen compared with trials when the target outside the RF is chosen. We therefore analyzed firing based on whether the selected target was in the RF (InRF trials) or out of the RF (OutRF trials; **Fig 1E,F** for LRS task. **Supp Fig 1-3** for ODR task). The VM neurons responded significantly more on InRF trials compared to OutRF trials (p=4.2 x 10^-3^, Wilcoxon rank-sum test, epoch=50 to 200 ms). Movement neurons also responded significantly more on InRF trials compared with OutRF trials, during movement but not immediately after target onset (p=5.7 X 10^-5^, Wilcoxon rank-sum test, epoch=150 to 250 ms). In contrast, visual neurons responded similarly for InRF and OutRF trials (p=0.61, Wilcoxon rank-sum test, epoch=0 to 100 ms).

One concern is that firing rate differences between InRF and OutRF conditions may reflect potential differences in the stimulus or reward alone. We therefore compared the firing rates for InRF and OutRF conditions on non-conflict trials in which only luminance changes or only reward changes (**Supp Fig 5**). VM neurons responded significantly more on InRF trials compared to OutRF trials, both when Bright target was selected (**Supp. Fig 5A**, Luminance-only trials, p=5.8 x 10^-4^, Wilcoxon rank-sum test, epoch=50 to 200 ms) and Rich target was selected (**Supp. Fig 5B**, Reward-only trials, p=4.5 x 10^-3^, Wilcoxon rank-sum test, epoch=50 to 200 ms). Movement neurons also responded significantly more on InRF trials compared with OutRF trials, during movement but not immediately after target onset (Luminance-only trials: p=1.1 x 10^-3^, Reward-only trials: p=3.8 x 10^-5^, Wilcoxon rank-sum test, epoch=150 to 250 ms). In contrast, visual neurons responded similarly for InRF and OutRF trials on reward-only trials (Reward-only trials, p=0.62 Wilcoxon rank-sum test, epoch=0 to 100 ms), but responded significantly more on InRF, luminance-only trials (**Supp Fig 5**, Luminance-only trials,p=0.03 Wilcoxon rank-sum test, epoch=0 to 100 ms). VM neuron responses are not simply due to differences in the stimulus or reward alone and are selective immediately after target onset. Visual neuron responses are not necessarily due to attention because they are only selective on luminance-only non-conflict trials. Movement neuron responses need not be due to attention because they are selective during the response and not immediately after target onset. These controls show that VM neurons play a more direct role in attentional selection than visual and movement neurons.

**Figure 5:**
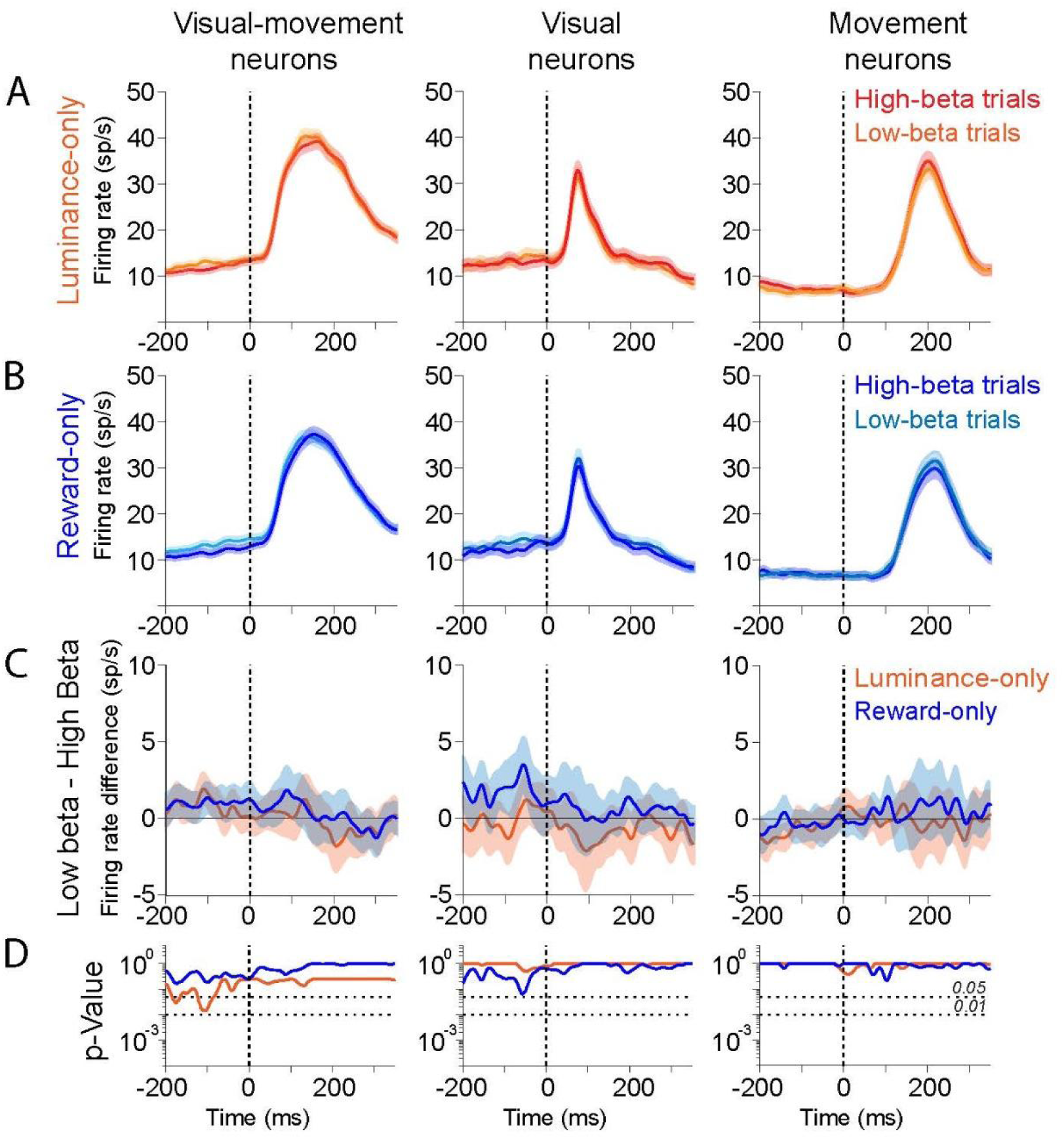
Beta-bursts do not modulate exogenous selection when not in conflict with endogenous selection. **(A)** Population average firing rates of three groups of neurons on luminance-varying trials. Luminance-only trials are non-conflict trials when one target is Bright and the other Dim and the reward values are the same. Mean +/- s.e.m. are shown for InRF trails when the Bright target is selected in the RF. Dotted lines denote target onset. **(B)** Same as A, but for reward-only trials. Reward-only trials are non-conflict trials when one target is Rich and the other Poor and luminance values are the same. Mean +/- s.e.m. are shown for InRF trails when the Rich target is selected in the RF. **(C)** Difference in firing rates for pre-target low beta-bursts (LB) and high beta-bursts (HB). Mean +/- s.e.m. are shown for exogenous (red) and endogenous (blue) selection. **(D)** Permutation test p-values under a null hypothesis that there is no difference in firing rates for high-beta and low-beta trials for exogenous selection (red) and endogenous selection (blue).

### Visual-movement neuron spiking reflects attentional selection

The LRS task reveals attentional selection by independently manipulating luminance and reward contingencies associated with two targets to yield two major sets of trials - congruent trials and conflict trials (**Fig. 2A**). On **congruent trials**, luminance and reward values are both high for one target (**Rich-Bright**) and are both low for the other target (**Poor-Dim**). Since sensory and reward drives are congruent, we expected that both favored selection of the same target regardless of saccade RT. This was observed in choice behavior. Each monkey had a strong preference for selecting the Rich-Bright target on congruent trials (M1: p=86%; M2: p=72%).

On conflict trials, one target had high-reward and low-luminance (**Rich-Dim**) and the other target had low-reward and high-luminance (**Poor-Bright**). Consequently, sensory and reward drives favored different targets. We hypothesized that LRS task performance on conflict trials may reveal endogenous, reward-driven selection or an exogenous, stimulus-driven selection (**Fig 2A**). We specifically predicted that RT should be longer on conflict trials when endogenous selection is expressed and the Rich-Dim target is chosen not the Poor-Bright target, *endo-conflict trials*, compared with conflict trials when exogenous selection is expressed and the Poor-Bright target is chosen, *exo-conflict trials*. RTs were significantly greater for endo-conflict trials compared to exo-conflict trials (M1: Endo-conflict RT=191+/-29ms, Exo-conflict RT=169+/-26 ms; p=5.3x10^-26^, M2: Endo-conflict RT=192+/-28ms, Exo-conflict RT=185+/-36 ms; p=8.2x10^-21^, Wilcoxon rank-sum test, mean +/- sem). Consequently, behavior on conflict trials revealed whether endogenous selection or exogenous selection was expressed trial-by-trial.

How behavioral choices varied with RT further support the importance of conflict trials in revealing exogenous and endogenous attentional selection processes. On conflict trials, shorter RTs reflected exogenous selection whereas longer RTs choices reflected endogenous selection (**Supp Fig 6**). In comparison, on luminance-only non-conflict trials shorter RTs reflected exogenous-driven selection. However, for longer RTs, the choice probability of selecting the bright target approached chance (**Supp Fig 6**). Similarly, on reward-only non-conflict trials longer RTs reflected endogenous-driven selection, however, for shorter RTs probability of selecting the rich target approached chance (**Supp Fig 6)**. Therefore, we hypothesized that conflict trials but not non-conflict trials reveal the mechanisms of exogenous and endogenous attentional selection.

**Figure 6:**
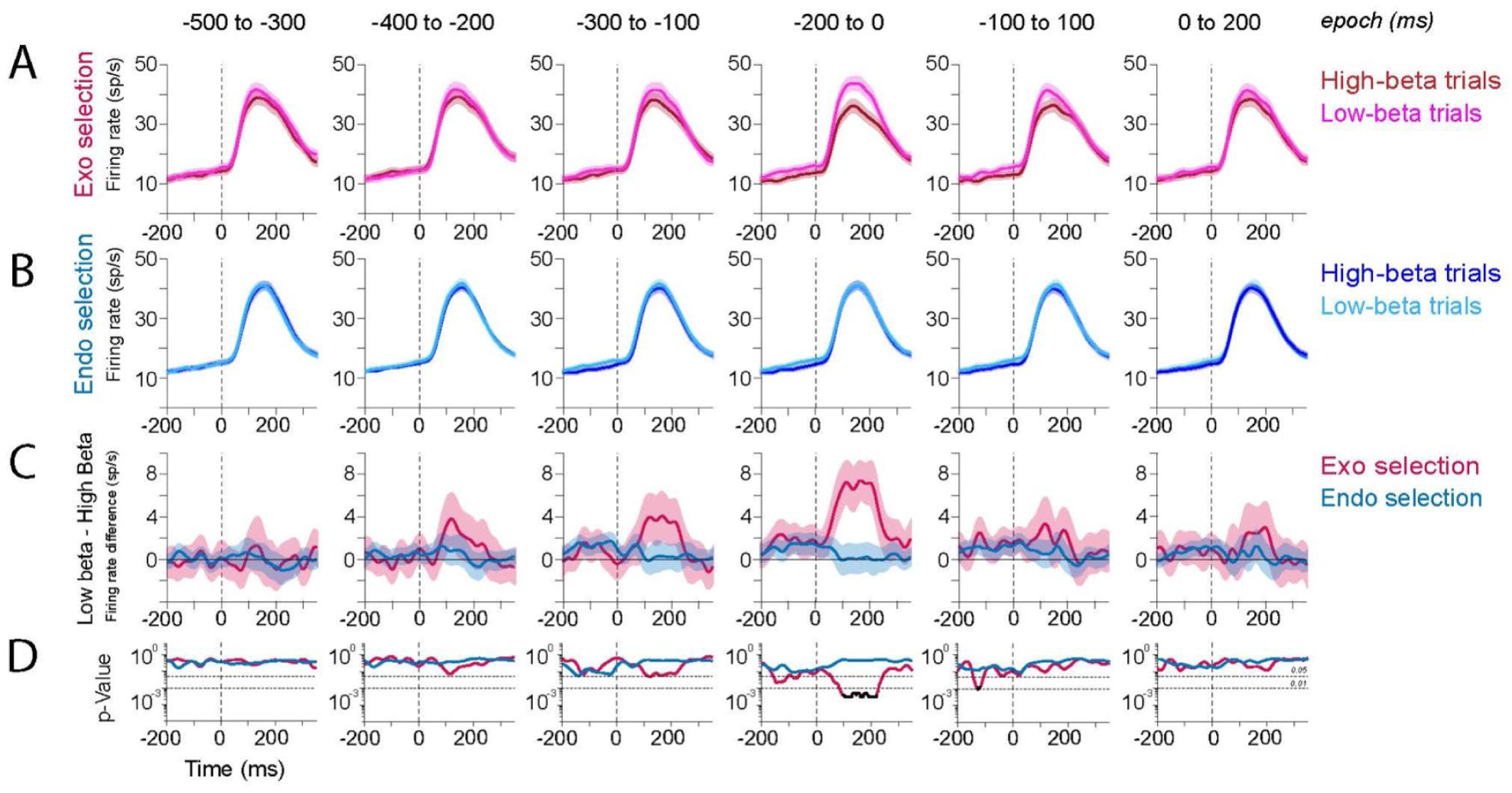
Beta-burst exogenous attentional modulation is transient in time. **(A)** VM neurons firing activity modulated by high (HB, darker traces) and low (LB lighter traces) beta-burst computed during six different time-windows. Mean +/- s.e.m. are shown for exogenous selection when the Poor-Bright target is selected in the RF. Dotted lines denote target onset. **(B)** Same as A, but for endogenous selection. Mean +/- s.e.m. are shown for InRF trials when the Rich-Dim target is selected in the RF. **(C)** Difference in firing rates for pre-target low beta-bursts (LB) and high beta-bursts (HB). Mean +/- s.e.m. are shown for exogenous (red) and endogenous (blue) selection. **(D)** Permutation test p-values under a null hypothesis that there is no difference in firing rates for high-beta and low-beta trials for exogenous selection (red) and endogenous selection (blue). FDR corrected for p-values for 0.01 alpha are shown in black.

We next investigated neural correlates of endogenous and exogenous attentional selection on conflict trials. We predicted that on exo-conflict trials when the Poor-Bright target is selected and the target is in the RF, **exo-InRF trials**, neuronal activity should differ from exo-conflict trials when the Poor-Bright target is selected and the target is out of the RF, **exo-OutRF trials** (**Fig 2B**). We also predicted that firing supporting endogenous attentional selection on endo-conflict trials when the Rich-Dim target in the RF is selected, **endo-InRF trials**, should differ from firing on endo-conflict trials when the Rich-Dim target out of the RF is selected, **endo-OutRF trials** (**Fig 2B**). Since exogenous attentional selection occurs earlier than endogenous attentional selection, we further predicted that neuronal selectivity on exo-conflict trials should occur earlier than on endo-conflict trials.

Consistent with a role in attentional selection during conflict, we observed that selectivity of VM neurons on exo-conflict trials occurred earlier compared to on endo-conflict trials. **Fig 2C** shows the responses of an example VM neuron for endo-conflict and exo-conflict trials. As expected, the VM neuron responded more when the InRF target was selected compared to when the OutRF target was selected for both exo-conflict trials and endo-conflict trials. Interestingly, VM neuron firing on inRF trials differed from OutRF trials substantially earlier on exo-conflict trials compared to endo-conflict trials. After the target onset, VM neuron firing rate during exogenous selection separated ∼50 ms earlier than during endogenous selection (Exo ST = 49 ms, Endo ST = 116 ms. **Fig 2D,E, Methods**, **Supp Fig 7**). Therefore, VM neurons process both exogenous and endogenous attentional selection during conflict.

**Figure 7:**
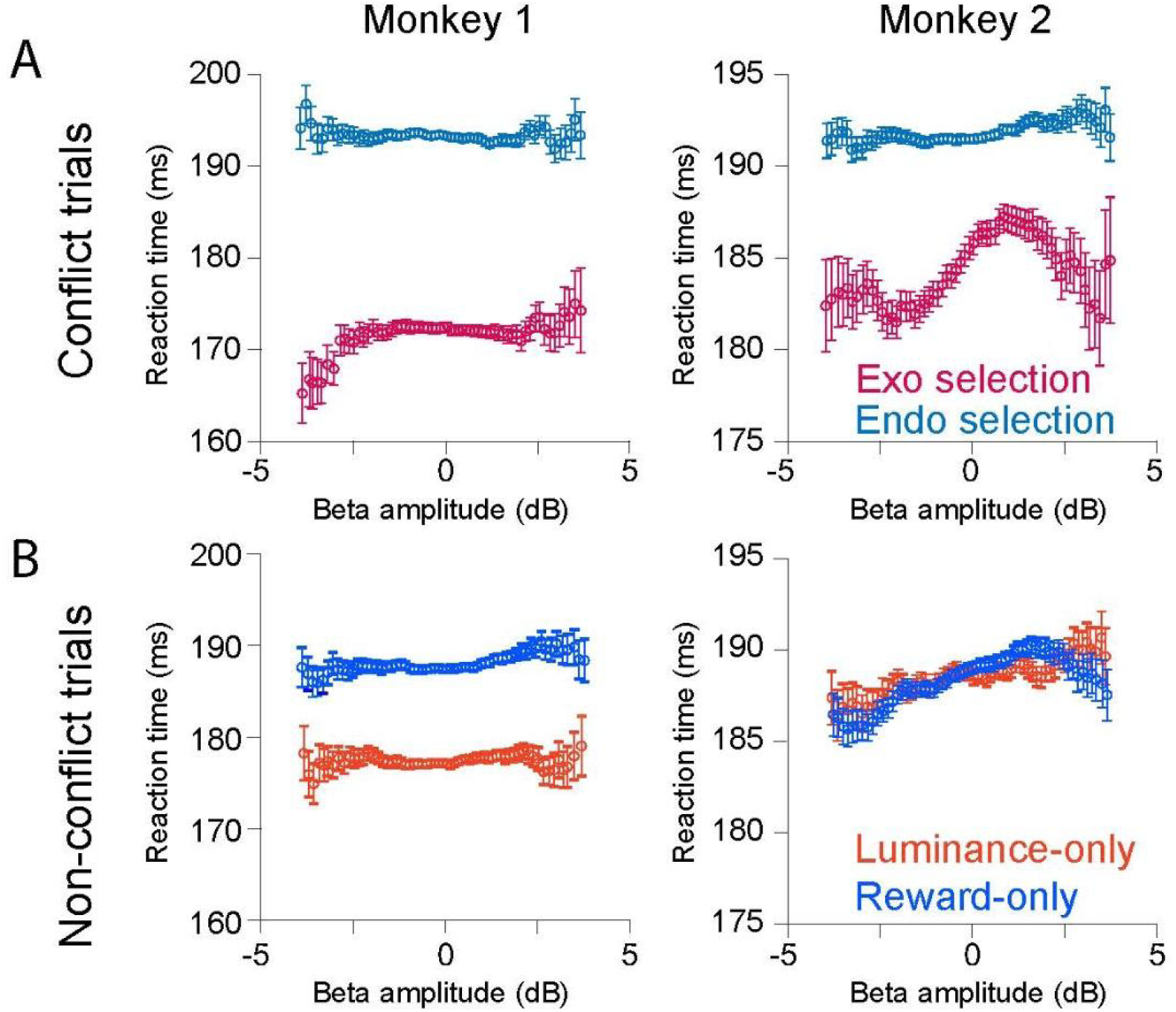
Beta-bursts selectively modulate exogenous selection reaction times conflict. **(A)** Saccade reaction times on conflict trials as a function of pre-target beta burst amplitude for Monkey 1 and Monkey 2. Exogenous selection choices are shown in red and endogenous selection choices are shown in blue. **(B)** Same as A, but for luminance-only and reward-only non-conflict trials.

The firing rates of visual neurons for InRF and OutRF conditions significantly differed on exo-conflict trials but not in endo-conflict trials (p<0.01, permutation test; Exo-ST=61 ms. **Fig 2E**). Consequently, visual neuron activity likely reflects processes related to exogenous selection alone and not conflict with endogenous selection. Movement neurons, on the other hand, showed elevated responses on InRF trials compared to OutRF trials for both exo-conflict and endo-conflict trials (**Fig 2E)**. But movement neuron firing rates for two conditions separated at a similar time after the target onset (p<0.01, permutation test, Exo ST = 153ms, Endo ST = 156ms, **Supp Fig 7**). This means that movement neuron activity does not reflect the conflict between endogenous and exogenous attentional selection. Movement neuron activity likely reflects processes which occur after attentional selection such as response preparation and the subsequent movement. Therefore, VM neuron firing and not visual or movement neuron firing specifically reflects endogenous and exogenous attentional dynamics during conflict.

### LPFC neuron spiking activity contains beta-frequency bursts

We next investigated the role of neuronal coherence in LPFC in the control of exogenous and endogenous attentional selection. In the pre-target period, LFP activity on individual electrodes displayed clear bursts of beta-frequency activity, 15-30 Hz, which we term beta-bursts **(Fig 3A**). Pre-target beta-bursts were clearly and reliably visible in LFP activity on individual trials. When present, beta-bursts tended to occur in the pre-target period and not after the target onset, and typically occurred for several hundred milliseconds.

For each trial, we estimated the amplitude of pre-target beta-bursts at a single site from 200 ms before target onset until target onset, a duration long enough to sample several cycles of activity at the beta-frequency. Beta-burst amplitude varied significantly from trial-to-trial, by almost a factor of 100 at the example site (**Fig 3B**). At each recording site, we tested whether beta bursts are specifically present in the 200ms prior to target onset against the null hypothesis of activity at other times during the trial using a permutation test. Across the population, beta bursts were reliably present across LPFC recording locations in each animal (M1: 1108 out of 1152 sites; M2: 1299 out of 1344, 96% of electrodes). Inspecting example trials for trials with low beta-burst amplitude revealed beta-bursts were effectively absent on these trials (see **Fig 3A** lower traces). We therefore grouped the trials with the highest ∼33% and lowest ∼33% beta-burst activity to yield high-beta (HB) trials and low-beta (LB) trials.

We first sought to assess whether beta-bursts in LFP activity could reflect a local source in LPFC. To help answer this question, we looked for evidence of coherent activity in the spiking activity of 409 single units in LPFC (M1: N=179. M2: N=230) by correlating spiking with nearby LFP activity (within approx. 1.5 mm) using spike-field coherence (SFC, **Fig 3C**). During the pre-target period, of the 409 neurons, 176 neurons significantly fired spikes at times predicted by nearby LFP activity in the beta-frequency range (15-35 Hz) (p<0.05 cluster corrected, permutation test; M1: N=59 and M2: N=117, **Supp. Fig 8**). This suggests that beta-burst LFP activity involves LPFC neuron firing and is not simply due to activity propagating from other regions that do not necessarily involve LPFC neuron firing. Interestingly, SFC amplitude in LPFC was greatest for activity in the beta-frequency range, compared with frequencies greater than 35 Hz. The number of LPFC neurons that fired coherently in the gamma (40-70 Hz) frequency range was not significant (<5 %, **Supp.Fig 8**).

**Figure 8:**
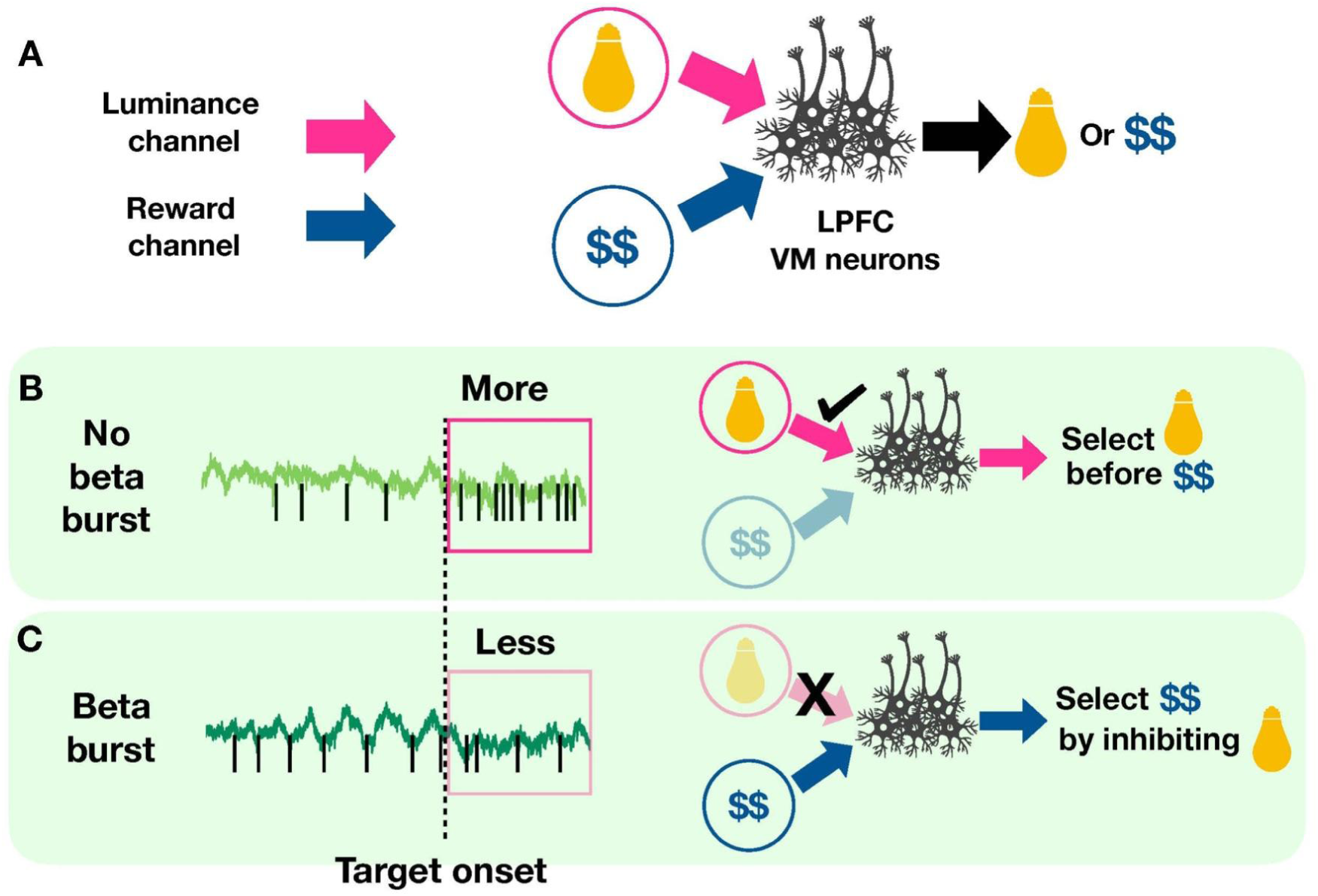
Channel modulation hypothesis supports attentional selection. **(A)** LPFC VM neurons receive luminance and reward information from two distinct communication channels, which compete to guide behavior. Exogenous selection of ‘Bright’ and endogenous selection of ‘Rich’ target depends on inhibitory modulation of the luminance channel. **(B)** In absence of beta-bursts, the luminance channel is open, communicating sensory-driven salient information earlier than goal-driven information. Information in the luminance channel drives the exogenous selection of the ‘Bright’ target. **(C)** In presence of beta-bursts, the luminance channel is close, inhibiting communication of sensory information. Information in the reward channel drives the endogenous selection of the ‘Rich’’ target.

Trial-to-trial variability in beta-burst amplitude may reflect trial-to-trial changes in the timing of spiking activity across the population of LPFC neurons. If so, spiking during HB trials should display greater coherence than spiking during LB trials. Furthermore, the dependence of neural coherence on beta-burst events should specifically be observed in the neurons that participate in the coherent activity. Neurons that do not participate, firing spikes at times that cannot be predicted by beta-frequency neural activity, should not show differences in coherence with beta-burst events. To test this, we estimated SFC immediately before target onset (see **Methods**) separately for the HB and LB trials for coherent and not-coherent neurons (**Fig 3D,E**). Consistent with a strong relationship between spiking and beta-burst events, SFC was significantly stronger during HB trials than LB trials for coherent neurons but not for not-coherent neurons (p=9.3 x 10^-27^ Wilcoxon rank-sum test). Importantly, the effect in coherent neurons was not simply due to the increase in LFP power during beta-bursts because SFC for not-coherent neurons was similarly insignificant during beta-bursts (p=0.1, Wilcoxon rank-sum test). These results demonstrate that when high-amplitude beta-bursts occur during the pre-target period they reflect increased coherent spiking in LPFC neurons.

SFC for VM, visual and movement neurons (**Supp Fig 9**) further show that beta-burst related coherence during the baseline is observed for the coherent neurons but not for the not-coherent neurons. Interestingly, the effect was weakest in the movement neurons, but comparable in the VM neurons as well as the visual neurons (VM neurons: coherent N=62, p=1.7 x 10^-15^, not-coherent N=77, p=0.23; Visual neurons: coherent N=24, p=1.2 x 10^-4^, not-coherent N=33, p=0.54; Movement neurons: coherent N=17, p=0.02, not-coherent N=48, p=0.9). This indicates that pre-target beta-bursts processes may specifically be involved in selection processes driven by visual input.

### Beta-bursts selectively modulate exogenous attentional selection during conflict

We next asked whether beta-bursts modulate attentional selection in general, or modulate either exogenous or endogenous attentional selection. Since the LRS task conflict trials dissociate endogenous and exogenous selection, we used these trials to test the relationship between beta-bursts and neuronal mechanisms of attentional selection, and whether beta-bursts exhibit specificity for endogenous or exogenous selection. On LRS conflict trials, selecting the Poor-Bright target in the presence of Rich-Dim target expressed exogenous selection. Similarly, selecting the Rich-Dim target in the presence of the Poor-Dim target expressed endogenous selection. We focused on LRS conflict trials for which the choices were made into the response field of each neuron under study.

We examined three hypotheses. First, since the firing rate of VM neurons reflects both endogenous and exogenous attentional selection, if LPFC beta-bursts modulate attentional selection, we specifically predicted that the rate of VM neuron firing would differ when beta-burst amplitude was high compared to when beta-burst amplitude was low. Second, since visual and movement neuron activity do not reflect attentional selection, if beta-bursts mediate control of attentional selection, we also predicted that the firing rate of these neurons should not differ on HB and LB trials. Finally, we predicted that if beta-bursts do not modulate selective attention in general and modulate either endogenous or exogenous selection, the relationship between beta-bursts and VM neuron firing rate should be present for either endogenous or exogenous selection trials and not both sets of trials.

Consistent with a role in attentional selection, we observed that VM neuron firing rate on InRF conflict trials involving exogenous selection significantly differed when pre-target beta-burst amplitude was high compared to when the amplitude was low (**Fig 4A**, InRF. VM-exo: p<0.01, permutation test). As expected, visual and movement neuron firing did not differ between HB and LB trials during exogenous selection (InRF conflict trials. Visual-exo: p>0.01, permutation test). This demonstrates that beta-bursts in LPFC can modulate attentional selection and do not modulate LPFC firing rates more generally.

Importantly, beta-bursts selectively modulated exogenous selection and LPFC neuron firing in the response field for HB and LB trials. Beta-bursts did not significantly modulate firing activity during endogenous selection and this was true for all three classes of neuronal response (InRF conflict trials. VM-endo: p>0.01; Visual-endo: p>0.01; Movement-endo: p>0.01, permutation test, **Supp Fig 10**). To further examine the nature of this beta-bursts modulation of attentional selection, we compared the VM neurons firing activity across all trials with choices into the response field against HB and LB trials (**Supp Fig 11**). On exo-HB trials, VM neurons firing rate was lower compared with the average firing rate, and on exo-LB trials VM neurons firing rate was higher compared with the average firing rate. This was not observed for endogenous selection trials (**Supp Fig 11**). These results demonstrate that an increase in pre-target beta activity suppresses luminance processing and a reduction in pre-target beta facilitates luminance processing selectively on exogenous selection trials.

It is possible that beta-bursts alter firing out of the response field and not into the response field. However, neuronal firing for VM, visual and movement neurons did not differ for exogenous and endogenous selection when beta-burst amplitude was high compared to low and the target outside the RF was chosen (OutRF: VM-exo: p>0.01. VM-endo: p>0.01. Visual-exo: p>0.01. Visual-endo: p>0.01. Movement-exo: p>0.01. Movement-endo: p>0.01, permutation test, **Supp Fig 12**). This suggests that beta-bursts modulate VM neurons firing rates for selection into the response field. Since LPFC neurons showed contralateral as well as ipsilateral response fields to the recording site, it is interesting to examine whether beta-burst modulation effects are different for contralateral-RF and ipsilateral-RF VM neurons. Majority of the VM neurons showed contralateral RFs (N=97) and reflected significant beta-bursts modulation effect on exogenous selection trials (**Supp Fig 13DE**). This was not observed for endogenous selection trials. VM neurons with ipsilateral RFs were smaller in number (N=25) and also reflected beta-burst modulation effects on exogenous selection, however the results were not statistically significant because of low sample size (**Supp Fig 13BC**). These results demonstrate that the beta-bursts modulation effect is specific to exogenous selection into the response field irrespective of the location of RF in the visual field.

One concern is that beta-burst modulation effect could be due to the grouping criterion used for selecting high-beta and low-beta trials. To examine whether the beta-bursts effect is specific to 33 % grouping, we selected high-beta and low-beta trials based on 50%, 40%, 25%, 15% and 10% grouping and examined beta-bursts effect on attentional selection (**Supp Fig 14B**). Beta-bursts modulation effect on exogenous selection was not selective to 33% grouping and was observed significantly for other percentiles as well (**Supp Fig 14B,C**). Importantly, beta-bursts grouping based on 50% grouping yielded similar results to 33% grouping.

It could be possible that the other frequency ranges in the pre-target period also reflect post-target modulation of VM neurons firing activity.. To test this we varied the Lfp frequencies range at every 5 Hz and examined VM neurons firing activity for attentional selection (**Supp Fig 15**). Exogenous effects were strongest in the 15-30 Hz frequency range and were specific to beta frequency ranges (**Supp Fig 15C**). Further, VM neurons firing rate were not modulated based on pre-target alpha (8-13 Hz) and gamma (40-70 Hz) activity (**Supp Fig 16**). These results demonstrate that LFP activity in the beta frequency range (15-30 Hz) specifically modulate neuronal mechanisms of exogenous attentional selection and do not modulate neuronal mechanisms of attentional selection more generally.

We next asked whether beta-bursts modulation effects alter the timing of exogenous selection processes on high-beta and low-beta trials. Since higher-amplitude beta-bursts suppress the post-target VM neurons firing (**Supp Fig 11**), we predicted that the timing of exogenous attentional selection processes would be delayed on high-beta trials compared to low-beta trials. Consistent with the suppression of VM neurons firing rate, higher-amplitude beta-bursts suppressed the exogenous selection process in time. Selection time on high-beta trials was 10 ms later compared to low-beta trials (exoHB ST = 54 ms, exo LB ST = 44 ms, p=0.03 permutation test, **Supp Fig 17 B,C**). This was not observed for endogenous selection (endoHB ST = 121 ms, endoLB ST = 117 ms, p=0.26, permutation test, **Supp Fig 17 E,F**). The difference in beta-burst amplitude values for high-beta exogenous and endogenous trials further supported the differences for exogenous and endogenous selection (p=0.02, Wilcoxon rank-sum test, **Supp Fig 18**). These results demonstrate that beta-burst activity modulates both the strength and timing of exogenous attentional selection processes.

### Beta bursts do not modulate exogenous selection in the absence of conflict with endogenous selection

On LRS conflict trials, exogenous and endogenous attentional selection processes are in conflict and compete. Choosing the Poor-Bright target in the presence of the Rich-Dim target expresses exogenous selection. We find that pre-target beta-bursts selectively inhibit the rate of VM neuron firing during exogenous selection on conflict trials. However, it remains unclear whether beta-bursts inhibit exogenous selection in general or reflect an active cognitive mechanism that is recruited during conflict.

Therefore, to further investigate beta-burst-related modulation of exogenous selection, we analyzed luminance-only non-conflict trials when luminance of the two targets differed, **Bright** and **Dim**, and the reward contingencies were similar across blocks (see **Methods**, **Fig 1Cb**). In these trials, there was no conflict present and selection for fast RTs was predominantly guided by the exogenous-selection processes (**Supp Fig 6**). We predicted that if beta-bursts inhibit exogenous selection in general, then VM neuron firing rate on InRF trials should differ on trials when beta-burst amplitude is high compared to when beta-burst amplitude is low. Alternatively, if beta-bursts specifically inhibit exogenous selection when there is conflict with endogenous selection, the rate of VM neuron firing should not differ on HB and LB trials.

Unlike during LRS conflict trials, the rate of VM neurons firing did not significantly differ for HB and LB trials when the Bright target was selected in the presence of the Dim target and the reward contingencies were the same (**Fig 5A**, InRF. VM-exo: p>0.01, permutation test and **Supp Fig 19**). Consistent with LRS conflict trials, visual and movement neuron firing did not differ between HB and LB trials on these trials (InRF. Visual-exo: p>0.01; Movement-exo: p>0.01, permutation test). This demonstrates that pre-target beta-bursts in LPFC specifically inhibits exogenous selection when in conflict with endogenous selection and does not modulate exogenous selection in general.

We also analyzed reward-only non-conflict trials when reward contingencies of the two targets differed, **Rich** and **Poor**, and the relative luminance of two targets were similar across blocks. The results confirmed that pre-target beta bursts did not modulate LPFC neuron firing rate in the absence of conflict between exogeneous and endogeneous selection (**Fig 5B**, InRF trials. VM-endo: p>0.01; Visual-endo: p>0.01; Movement-endo: p>0.01, permutation test).

### Pre-target beta-burst exogenous attentional modulation is transient in time

Pre-target beta-bursts (-200 ms to 0 ms, where 0 is target onset) selectively inhibit the neuronal mechanisms of exogenous attentional selection and not the endogenous attentional selection. Since exogenous selection occurs earlier in time (49 ms) compared to endogenous selection (116 ms), one concern is that the results may simply be due to the proximity of beta-bursts in time to exogenous-selection mechanisms. If so, beta-bursts that occur later in time, and hence closer to the time of endogenous selection, may instead modulate endogenous-selection not exogenous-selection. To address this concern, we analyzed beta-bursts during six time epochs and studied VM neuron firing patterns for InRF trials involving endogenous-selection when beta-burst amplitude was high compared to when the amplitude was low (**Fig 6B**). VM neuron firing rates on InRF trials were not significantly different for high-beta and low-beta trials during the [-100 100] epochs (**Fig. 6C,D**, Endo [-100 100]: p>0.01, permutation Test).

Examining the time course of beta-burst related modulation also revealed that early beta-bursts do not tend to modulate exogenous-attentional selection (**Fig 6A**). The strongest modulation of VM neuron firing was observed for beta-bursts that occurred immediately before target onset **Fig. 6C.**

### Pre-target beta-bursts modulate exogenous selection reaction times

If the selective modulation of VM neuron firing with coherent-beta activity reflects attentional selection, then the modulatory effect of beta activity on conflict trials should be present during exogenous choice behavior more than during endogenous choice behavior. Since behavioral RTs reflect the underlying mechanism of attentional selection, we specifically predicted that RTs should vary trial-by-trial with coherent beta-activity on exo-conflict trials more than on endo-conflict trials. For each monkey, changes in coherent-beta activity were associated with changes in RTs on exogenous choice conflict trials more than on endogenous choice conflict trials (**Fig 7A**, see **Methods**). For the exo-conflict group of trials, the reaction times significantly differed with coherent-beta activity (M1: RT Range = 4.65%. p = 0.043. M2: RT range = 2.97%. p = 1x10^-3^. Permutation test; RT range refers to range (max RT - min RT) of variation of RT with beta values). For the endo-conflict group of trials, RTs did not significantly differ with coherent-beta activity (M1: RT range = 1.25%. p = 0.91. M2: RT range = 1.12%. p = 0.53. Permutation test). Finally, since VM neuron firing effects are not present on non-conflict trials, we also predict that the relationship between beta-activity and RTs should not be present on non-conflict trials. Consistent with VM neuron firing effects, pre-target coherent-beta activity and RTs on non-conflict trials did not significantly differ when sorting by either group of trials (**Fig 7B. Luminance-only:** M1: RT range = 1.21%. p = 0.928. M2: RT range = 3.62%. p = 1. **Reward-only:** M1: RT range = 2.63%. p = 0.156. M2: RT range = 2.41%. p = 1. Permutation test). Therefore, the role of coherent-beta activity in attentional selection is specifically present during conflict, is consistent with the pattern of results observed for the VM neurons and consequently may mediate top-down control of attentional selection by modulating sensory, cue-driven responses in the VM neuron subpopulation.

## Discussion

Here, we make two specific contributions that demonstrate a role for beta-frequency neural coherence in attentional selection through inhibitory mechanisms (**Fig 8**). We propose that attentional selection involves filtering of luminance and reward channels that communicate information to visual-movement neurons in the lateral prefrontal cortex in order to select a response (**Fig 8A**). When beta bursts are not present, target onset drives LPFC to select information in the luminance channel before information the reward channel is available (**Fig 8B**). When beta bursts are present, information in the luminance channel is inhibited and the response is selected based on information in the reward channel (**Fig 8C**).

We then demonstrate that coherent neuronal activity in the beta frequency range (15-30 Hz) selectively modulates exogenous selection by suppressing the luminance channel providing salient sensory information. Beta activity observed in the pre-target period is associated with the inhibited post-target, sensory-driven firing by LPFC neurons when selection is driven by exogenous attention but not by endogenous attention. Consequently, our results are consistent with the top-down control view of attentional selection. According to the top-down control view, the selection of task-relevant endogenous targets relies on mechanisms of multiregional communication that limit the influence of sensory inputs (Anderson and Weaver, 2009; Miller and Cohen, 2001; Womelsdorf and Everling, 2015). Since top-down control mechanisms operate under the knowledge of task-relevance (Miller and Cohen, 2001), the beta-activity effect was observed on conflict trials but not on non-conflict trials. On conflict trials selection of the task-relevant target yielded high reward whereas on non-conflict trials the task did not prioritize one target with another based on reward-value. As the role played by coherent-beta activity is to modulate information flow due to sensory inputs, our work provides new evidence for how coherent-beta activity in LPFC could mediate the top-down control of attentional selection.

We also show how coherent-beta-activity could bias the mechanisms of attentional selection in LPFC by influencing the flow of sensory information during target selection. We show that a subgroup of LPFC neurons, termed visual-movement (VM) neurons and not visual and movement neurons, encodes both exogenous and endogenous attentional selection. The timescales underlying exogenous and endogenous selection have been a major focus of behavioral work which has shown that reaction times are typically ∼30 ms faster for exogenous selection (Awh et al., 2006; Carrasco, 2011; Corbetta and Shulman, 2002; Markowitz et al., 2011; Theeuwes, 2010). Here, we go further and measure the timescales of exogenous and endogenous selection by analyzing the spiking patterns of populations of individual LPFC neurons. Consistent with previous recordings in LPFC (Buschman and Miller, 2007), we find that VM neuron spiking activity in response to target onset encoded exogenous selection ∼50 ms before endogenous selection. This difference in timing means that coherent-beta-activity in PFC can have a substantial influence on the direction of sensory information flow and bias the selection of relevant targets in the presence of irrelevant distractors.

In the following, we discuss the mechanisms of top-down attentional control and how coherent-beta activity may support the selection of task-relevant targets.

### Top-down attentional control mechanisms are mediated by coherent-beta activity

In LPFC, beta activity reflects exogenous and endogenous attentional selection processes (Bastos et al., 2015; Buschman and Miller, 2007; Buschman et al., 2012; Fiebelkorn and Kastner, 2021). The emergence of LPFC beta activity before selection during different goal-defining tasks further suggests a role for beta activity in the top-down control of attention selection (Bastos et al., 2015; Buschman et al., 2012; Womelsdorf and Everling, 2015). Here, we more closely examine the strong relationship between spiking and coherent beta activity in LPFC immediately before presenting relevant and irrelevant targets to reveal mechanisms of top-down attentional control. The central aspect of top-down control is inhibition with knowledge of what needs to be controlled, i.e. relevance (Miller and Cohen, 2001). We show that LPFC beta activity is associated with the inhibition of LPFC neural firing during exogenous selection and not endogenous selection, and so is grounded in task-relevance. Importantly, LPFC-beta-activity-mediated selective inhibition was only observed in presence of conflict i.e, when sensory and reward drive each favored the selection of different targets (**Fig 4**). In absence of conflict, when sensory information was absent, LPFC firing rates were not modulated with beta activity (**Fig 5**).

On conflict trials, reward-drive favored the selection of the task-relevant target whereas, on non-conflict trials, absence of reward-drive diminished the task relevance of one target over other. Therefore, we propose that LPFC performs top-down control of attentional selection by deploying beta-frequency coherent neural activity to selectively limit or bias the flow of sensory information specifically when conflicting information drives target selection.

The posterior parietal cortices also process exogenous sensory information (Buschman and Miller, 2007; Chen et al., 2020; Gottlieb et al., 1998; Suzuki and Gottlieb, 2013). Consequently, the LPFC coherent-beta network that selectively inhibits sensory information likely operates across frontal-parietal projections. Indeed, frontal and parietal areas both reflect coherent-beta activity indexing stimulus selection in attention and working memory (Buschman and Miller, 2007; Fiebelkorn and Kastner, 2021; Salazar et al., 2012). LPFC may selectively inhibit PPC information flow through a long range beta network. If so, prefrontal areas need to generate a sufficiently reliable and impactful neural stimulus to influence posterior parietal areas. The firing of bursts as compared to single isolated spikes offer a candidate mechanism (Ardid et al., 2015; Lisman, 1997). For example, long range beta-burst synchronization between anterior cingulate cortex and LPFC exists during selective attention (Womelsdorf et al., 2014). Our observations of pre-target beta-bursts highlight a potential mechanistic role in how information is routed through PPC for the top-down control of attentional selection.

### LRS task reveals the timescale of exogenous and endogenous selection mechanisms

Our LRS task design was also necessary to reveal the time-course of attentional selection mechanisms in LPFC activity. In particular, the use of a non-cue binary choice task design where selection was made immediately after the target onset without any delay, and the use of spatial randomization of both target locations trial-by-trial revealed the timescales of different forms of attentional selection. Previously-used behavioral task designs have often manipulated spatial attention in a delayed design by presenting an attentional cue before the onset of a target (Bisley and Goldberg, 2003; Buschman and Miller, 2007; Fiebelkorn and Kastner, 2021; Fiebelkorn et al., 2018; Liu et al., 2009; Reynolds et al., 1999). In such paradigms, spatial attention is allocated to the cue location before exogenous or endogenous attention is recruited by the target. Previous work has also used task designs in which targets are presented at spatial locations in a predictable manner, which can generate spatial biases in behavior that may also be confounded with attentional selection mechanisms (Liu et al., 2009).

Our LRS task design was critical in dissociating exogenous and endogenous attentional selection mechanisms. Interestingly, however, previous work has shown that a non-salient target previously associated with reward may also capture attention (Anderson et al., 2011; Jahfari and Theeuwes, 2017), resulting in an interaction between salience-drive and involuntary-value-driven automatic attention. Whether coherent beta activity is implicated in the neural mechanisms of interactions between other forms of attention capture is an interesting direction for further work.

It could be argued that the difference in VM neurons firing modulation for exo and endo conditions is simply due to physical brightness of the target and not attentional selection (**Fig 2**). However, the luminance-only non-conflict trials in our task provide the necessary control for physical brightness effects. In these trials, physical brightness is the same as in the conflict trials but the recruitment of exogenous and endogenous attention differs (**Supp Fig 6**). Further, the physical brightness value of the selected target was not correlated to the pre-target beta value (**Supp Fig 20**). Therefore, the beta-modulation effects in our report are most consistent with an attentional effect.

### Dynamic interplay of exogenous and endogenous attentional selection

By dynamically shifting between more active and less active coherent states, high-beta and low-beta, our results show that VM neurons in the coherent-beta subnetwork may flexibly modulate multiregional communication across a sensory information channel that carries visual target information into the association cortices. We report behavioral effects in which the influence of coherent-beta-activity on saccade RTs is consistent with the effects observed in VM neuron firing. We find that high-beta activity selectively modulates response time specifically when making exogenous choices (**Fig 7**). Changes in coherent-beta activity were associated with changes in RTs on exogenous choice conflict trials more than on endogenous choice conflict trials. On trials when the choice was to the endogenous target, RTs were more similar across trials with beta bursts before target onset that differed in strength. This pattern of results mirrors that for the variations of VM neuron firing with pre-target beta activity across conflict and non-conflict trials. Thus, neural and behavioral results reinforce the flexible interplay between exogenous and endogenous selection which results from beta-activity mediated dynamic modulation of a sensory-driven information channel.

### Comparison with previous studies

Previous studies have associated beta activity with inhibition and reach movement initiation (Dean et al., 2012; Hagan et al., 2012; Pape and Siegel, 2016; Sanes and Donoghue, 1993). In the sensorimotor cortex, beta amplitude increases at rest and for stable postures and reduces during movement (Cassim et al., 2001; Kilavik et al., 2013; Schmidt et al., 2019). For example, Kilavik et al. showed increased beta in both pre-cue and pre-go epochs of reach movement tasks, with a temporary drop in beta amplitude after the cue (Kilavik et al., 2012). The post-cue suppression of beta-amplitude for movement planning and initiation may draw parallel to PFC beta observed before oculomotor selection in pre-target period. However, the detailed pattern of our results does not suggest that the PFC beta is related to the eye-movement itself. We only observe the beta modulation effects on trials involving conflict. If the results were due to movement suppression we would also observe them on the reward and luminance only varying trials (Fig 5). We do not observe the beta effects on the PFC neurons whose activity is most tied to the movement - the movement neurons. We observe the influence on neurons that have visual-movement responses (Fig 4). Furthermore, beta activity altered VM neurons firing for both ipsilateral and contralateral saccade selection and is not a lateralized motor effect (Supp Fig 13). Finally, we do not observe the effects on trials captured by endogenous attention and only on trials with luminance driven exogenous responses (Fig 4). These features of our results are not simply explained by movement suppression.

Prefrontal beta-activity observed in attention and working-memory tasks reflects information about the task-relevant rules that determine stimulus mapping to responses (Buschman and Miller, 2007; Buschman et al., 2012; Salazar et al., 2012). In these studies, the task-rule was to find a match for the sample either in object or space feature after a cue/delay period. In comparison, the LRS task utilized a non-cue binary choice task design to examine attentional selection mechanisms. Beta-bursts modulation effect in this report could be assigned to differences in involvement of beta-activity in the preparatory period and cue-triggered delay-period. However, we propose that it is the conflict between the two sources of information that influences the attentional selection process irrespective of the beta activity period - cue-triggered/preparatory beta. The LRS task design differed from typical attentional task designs in involving two competing sources of information during the target-period. The channel modulation top-down control model (**Fig 8**) suggests that beta-activity is involved in resolving conflict. Therefore, cue-period beta-activity may also influence the selection process in the presence of conflict.

Previous studies have suggested that LPFC neurons firing activity is modulated by reward-value (Kaping et al., 2011; Leon and Shadlen, 1999). We did not observe a value based modulation of VM neuron firing activity (**Supp. Fig. 21).** Interestingly, however, we did observe a firing rate modulation for movement neurons (**Supp. Fig. 21)**. Movement neuron firing activity was higher when the rich target was selected inRF compared to poor target selection inRF. Movement neuron activity during our task likely does not reflect endogenous attention processing and may instead reflect a form of reward expectancy. Note, however, the increased firing activity for the rich target is contrary to the Kaping et al. observation that shows an enhancement in LPFC activity when a low value target was selected over a high value target. This discrepancy could arise from our use of an immediate saccade unlike other work involving covert attentional cues.

In conclusion, we reveal the mechanisms of top-down control of attentional selection in LPFC involve the inhibition of luminance information to facilitate reward-guided behavior. We show that the dynamics of a population of VM neurons that fire coherently with beta activity may mediate top-down control of attentional selection, consistent with a role in inhibitory multiregional communication. We further show that coherent beta-activity selectively modulates exogenous responding compared with endogenous responding resulting in the flexible interplay between exogenous and endogenous selection necessary to resolve conflict.

## Acknowledgements

We would like to thank Gerardo Moreno for surgical assistance, Roch Comeau, Stephen Frey and Brian Hynes for custom modifications to the BrainSight system, and members of the Pesaran lab for helpful feedback. We thank Supratim Ray for insightful comments on an earlier version of this manuscript. This work was supported in part by NIH Ruth L. Kirschstein National Service Award F32-MH100884 from the National Institute of Mental Health (NIMH) (D.A.M.), a Swartz Fellowship in Theoretical Neurobiology (D.A.M.), NIH Training Grant T32-EY007158 (D.A.M.), R01 - NS104923 (B.P.) and MURI W911NF-16-1-0368 (B.P.). D.A.M and B.P designed and performed the experiments. A.D and B.P analyzed the data. A.D. and B.P. wrote the manuscript. The authors declare no competing interests.

## STAR Methods

### Experimental preparation

All surgical and animal care procedures were done in accordance with National Institute of Health guidelines and were approved by the New York University Animal Care and Use Committee. Two adult male rhesus monkeys (*Macaca mulatta*) participated in the experiments (Monkey 1, 9.5 kg and Monkey 2, 8.4 kg). Both animals had been previously used in other eye-movement experiments (Markowitz et al., 2011, 2015). Once trained on behavioral tasks, each animal was implanted with a low-profile recording chamber (Gray Matter Research, MT). The craniotomy was made over the right pre-arcuate cortex of each animal using image-guided stereotaxic surgical techniques (Brainsight, Rogue Research, Canada). A semichronic microelectrode array microdrive (SC32-1, Gray Matter Research, MT) was inserted into the recording chamber and sealed. The SC32-1 system has 32 microelectrodes, spaced 1.5 mm away (Fig 1D). The SC32-1 is a modular, replaceable system capable of independent bidirectional control of 32 microelectrodes.

### Behavioral experiments

#### Experimental hardware and software

Eye position was constantly monitored with an infrared optical eye tracking system sampling at 120 Hz (ISCAN). Visual stimuli were presented on an LCD screen (Dell Inc) placed 34 cm from the animal’s eyes. The visual stimuli were controlled via custom LabVIEW (National Instruments) software executed on a real-time embedded system (NI PXI-8184, National Instruments).

#### Experimental design

Each monkey first performed a **visually-guided oculomotor delayed response (ODR) task** to map the spatial response fields of neurons. Each monkey then performed the **luminance-reward-selection (LRS) task** to study the flexible control of attentional selection. Behavior and neural data was recorded across 39 (Monkey 1) and 42 (Monkey 2) experimental sessions.

#### ODR task

Each trial began with a visual fixation target presented at the center of the screen. Each animal maintained fixation for a variable 500-800 ms baseline period. After the baseline period, a red square appeared in the periphery to indicate target location of the saccade. There were eight possible iso-eccentric target locations spaced 10 deg around central fixation. Target location was randomized over trials so that animals could not predict where the cue would appear on any given trial. Each monkey maintained fixation for a variable 1000-1500 ms delay period. After the delay period, the central fixation square was extinguished, providing the Go signal for the animal to move his eyes to the target location. A fluid reward was awarded on successful completion of the trial. A trial was aborted if the animal failed to align his gaze within 2deg of the center of fixation or periphery target. On a given experimental session, on average M1 95 +/- 24 trials performed trials and M2 performed 248 +/- 32 ODR trials (mean +/- sd).

*LRS task:* Each trial again started with fixation at a visual target at the center of the screen for a variable 500-800 ms baseline period. After the baseline period, the center fixation target was extinguished, and two red targets (**T1 and T2**) were presented at random locations in the visual periphery at a 10 deg eccentricity from the central fixation. Two targets were constrained to be at least 90 deg apart on each trial. The randomized spatial location of targets controlled for the influence of spatial attention at the start of each trial. Onset of targets provided the animal Go signal to perform a saccade to one of the targets. Each animal was required to maintain a fixation of 300 ms at the chosen target, after which appropriate juice reward was delivered. Each trial lasted 890-1400 ms, and only one choice could be made per trial. A trial was aborted if the animal failed to align his gaze within 2deg of the center of fixation or choice targets. On a given experimental session, on average M1 performed 1276 +/- 348 and M2 performed 1677 +/- 139 (mean +/- sd) LRS trials.

**T1** and **T2** were two identical in size rectangular stimuli (3-to-1 aspect ratio) with different orientation (Fig 1B). T1 was oriented so that the long axis was vertical and T2 was oriented so that the long axis is horizontal. Long axis of each target subtended 2 deg of visual arc. Two targets were associated with different liquid reward values. Each animal was motivated to select the target associated with the highest value of liquid reward. Mean value of the liquid reward associated with each target was kept constant for blocks of 40-70 trials (Fig 1C). The block transition was unsignaled. Mean reward values varied between 0.04 ml/trial and 0.21 ml/trial. On each trial, a Gaussian-distributed variability (SD = 0.015 ml) was added to the value with each target. Variable reward values further increased animal’s uncertainty about the times of reward block transitions. Since the choice behavior around each reward block transition was more exploratory (**Supp. Fig 22**), we performed all the analysis after excluding the first 10 trials after the block transition. This ensured that the animals followed the reward contingencies.

On each trial, target luminance values were randomly assigned. T1 luminance was randomly assigned from a log-uniform distribution of values ranging from 0.01 to 12.15 cd/m^2^. The minimum luminance value was set above the psychophysical threshold for stimulus detection titrated during the ODR task. After the T1 luminance was assigned, the luminance of T2 was assigned such that mean luminance across both targets was 6 cd/m^2^. On each trial, target luminance values were assigned independently from the rewards associated with T1 and T2. Additionally, the randomized spatial locations of two targets ensured that the target location of the high-reward and low-luminance target could not be determined from the low-reward and high-luminance target.

Trial-by-trial independent manipulation of luminance and reward values randomly yielded either **congruent** or **conflict** set of trials. On a given experimental session, on average Monkey 1 performed 322 +/- 74 congruent trials and 317 +/- 81 conflict trials; Monkey 2 performed 392 +/- 33 congruent trials and 392 +/- 39 conflict trials (mean +/- sd).

On congruent trials, luminance and reward values were both high for one target (**Rich-Bright**) and were both low for the other target (**Poor-Dim**). Each monkey showed a strong preference for selecting **Rich-Bright** target compared to **Poor-Dim** target (M1: 84% total trials: 9881; M2: 72% total trials 15615: across 39 and 42 experimental sessions).

On conflict trials, however, one target had high-reward and low-luminance (**Rich-Dim**) and the other target had low-reward and high-luminance (**Poor-Bright**). Conflict trials, when endogenous selection was expressed and Rich-Dim target was selected were termed as **endo-conflict** trials (on average each monkey performed M1=211 +/- 62, M2= 301+/- 44 endo-conflict trials per experimental session, mean +/- sd). Similarly, conflict trials when exogenous selection was expressed and Poor-Bright was selected, were termed **exo** trials (on average each monkey performed M1=107 +/- 47, M2= 91 +/- 25 exo-conflict trials per experimental session, mean +/- sd). Each monkey followed rewards and showed preference for selecting **Rich-Dim** target compared to **Poor-Bright** target (M1: 68% total trials: 9751; M2: 77% total trials: 15652 trials, across 39 and 42 experimental sessions).

On each experimental session, on a subset of trials, the LRS task featured non-conflict **reward-only** and **luminance-only** trials. On reward-only trials, the luminance values of two targets were kept the same for blocks. On luminance-only trials, the average reward values associated with two targets were kept the same for blocks. On a given experimental session, on average Monkey 1 performed 220 +/- 98 reward-only trials and 294 +/- 120 luminance-only trials and Monkey 2 performed 245 +/- 29 reward-only trials and 337 +/- 57 luminance-only trials (mean +/- sd).

### Neurophysiological experiments

#### Recording protocol and data acquisition

Neural recordings were made with glass-coated tungsten electrodes (Alpha Omega, Israel) with impedance 0.7-1.5 M measured at 1 kHz (Bak Electronics, MD). Neural signals were preamplified (10 x gain; Multichannel Systems, Germany), amplified and digitized (16 bits at 30 kHz; NSpike, Harvard Instrumentation Lab), and continuously streamed to disk during the experiment (custom C and Matlab code). Neural recordings were referenced to a ground screw implanted in the left occipital lobe, with the tip of the screw just piercing through the dura mater.

In each animal, electrodes were advanced in each recording session to maximize the yield of isolated single units. Electrodes were advanced through a silastic membrane in the recording chamber, the dura mater and pia before entering the cortex. Each electrode was advanced sequentially in increments of 15 microns, 10 minutes apart to give the electrode time to settle in the tissue. Initial action potentials were recorded at a median depth of 3 mm (2.23 mm in M1; 3.04 mm in M2). Electrodes were gradually advanced across sessions (on average 34 µm/day in M1 and 100 µm/day in M2) until action potentials were no longer present, indicating passage into white matter. Neural recordings were made up to a median distance of 6 mm from their initial position.

Local field potential (LFP) activity was obtained offline by low-pass filtering the broadband raw recording at 300 Hz using a multitaper filter with a 1.5 ms time window. The low-pass filtered LFP activity was further downsampled to 1 kHz from 30 kHz. Multiunit activity (MUA) was obtained by high-pass filtering the raw recordings at 300 Hz and maintaining the original 30 kHz sampling rate. Single unit activity (SUA) was isolated by thresholding MUA activity at 3.5 standard deviations below the mean, performing a principal component analysis of putative spike waveforms, over-clustering these waveforms in PCA using k-means and then merging clusters based on visual inspection. Spike-sorting was performed for each recording session using custom Matlab code (Mathworks). Non-stationarity in recordings were accounted for by performing spike-sorting in 100 ms moving windows. Trials on which spike-clusters were not isolated were removed from further analysis.

#### Neuronal databases

We advanced electrodes to isolate and record 746 units (M1: 384; M2: 362 units) during the ODR task. Out of 746 units, we further selected 409 (M1: 179; M2: 230 units) single units that were responsive to the LRS task. We selected units with firing rates greater than 5 sp/s in 0 to 200 ms epoch after onset of targets for the LRS task.

Each neuron’s response-field (RF) was mapped using the ODR saccade task to eight possible target locations. LPFC neurons showed increased firing in response to target onset alone, saccadic eye movement alone or both target onset and saccadic eye movement (Fig 1). Therefore, we computed each neuron’s trial-averaged baseline subtracted firing rate in response to eight target locations around target onset and saccade onset (Target onset: baseline epoch = [-200 0ms], stimulus epoch = [0 100ms] and [75 200ms] where 0ms is targets onset; Saccade onset: baseline epoch = [-400 200ms], stimulus epoch = [-50 70ms] where 0ms is saccade onset). We used these epochs to accommodate the firing activity of visual, visual-movement (VM) and movement neurons (Supp Fig 1-3). Each neuron’s RF was estimated against the null hypothesis that there is no difference in response firing rate with respect to baseline, using a permutation test. The baseline-subtracted firing rate at each target’s location was compared with the null distribution. Null distribution was generated by shuffling firing rate across eight target locations 1000 times (p<0.05, permutation test). Since this procedure involves multiple comparisons, we corrected the p values by controlling for the false discovery rate (FDR, (Benjamini and Hochberg, 1995). Units with significant p-values either for target onset or saccade onset epochs were used for further analysis. Out of the 409 single units, we selected 216 neurons that showed an excitatory response inside the RF and had greater than 5 Hz firing rate either around target or saccade epoch (M1 = 122; M2 =139 neurons) .

The ODR task further revealed the firing patterns of different LPFC neurons. We classified each unit that had an excitatory RF response into visual, visual-movement (VM) and movement neurons based on their firing patterns around target onset and saccadic eye movement. The delay period of the ODR task separated the visual and saccade related neuronal activity and allowed us to examine each neuron’s firing patterns in response to target and saccade onset. Around target onset, visual and VM neurons showed an increase in firing activity and not movement neurons. Additionally, visual neurons reflected an increase in firing rate immediately after the target onset whereas VM neurons showed a delayed response (see **Supp Fig 1-2**, and **Fig 1**). Around saccade onset, VM and movement neurons showed an increase in firing activity and not visual neurons. Single unit responses at preferred target location were tested for selectivity around target onset and saccade onset through permutation testing. To classify between visual and VM neurons we compared each unit’s baseline-subtracted firing rate around target onset epochs (0 to 100 ms and 75 to 200 ms, where 0 ms is target onset). To classify between movement and VM neurons we compared each unit’s baseline-subtracted firing rate around saccade onset epoch (-50 to 70 ms where 0 ms is saccade onset). Units with significant p-values in target-onset (0 to 100 ms) epoch and not saccade-onset epoch were classified as visual neurons. Units with significant p-values in saccade-onset epoch and not target-onset epoch were classified as movement neurons. Units with significant p-values in both target-onset (75 to 200 ms) and saccade-onset epochs were classified as VM neurons. We further confirmed each unit’s classification label by visual inspection. Out of 261 units, N=139 (M1=54, M2=85) were VM neurons, N=57 (M1=27, M2=30) were visual neurons and N=65 (M1=41, M2=24) were movement neurons.

### Data analysis

#### LRS task selectivity

On the LRS task, the two targets were presented simultaneously. Therefore, on each trial, the location of both the targets with respect to a LPFC neuron’s RF was identified. For further analysis, we pooled the data across two monkeys to increase the statistical power. For each neuron, we selected the subset of trials on which one target was inside the RF and the other was outside the RF. Trials on which both the targets were inside the RF or both the targets were outside the RF were removed from further analysis. We examined each neuron’s selectivity to the LRS task based on the saccade response and the target properties. Trials on which saccade response was inside the RF were termed as InRF trials and trials on which saccade response was outside the RF were termed OutRF trials. Fig 1F shows the population data of 139 VM neurons across 36864 InRF trials and 36455 OutRF trials. Similar to the ODR task, VM neurons responded significantly more on trials when the InRF target was selected compared to trials when the OutRF target was selected (p=4.2 x 10^-3^, Wilcoxon rank-sum test, epoch=50 to 200 ms). Firing rate increased soon after target onset and extended through the saccade. Movement neurons (N=65) also responded significantly more on the InRF (N=16450) trials compared to OutRF (N=16020) trials (p=5.7 x 10^-5^, Wilcoxon rank-sum test, epoch=150 to 250 ms). Visual neurons (N=57) however, showed comparable firing rates for InRF (N=14416) and OutRF (N=15651) trials (p=0.61, Wilcoxon rank-sum test, epoch=0 to 100 ms). The results were similar if different time-windows around the peak-firing rates were used ([56 304], [0 182], and [139 301] for VM, movement and visual neurons. These time-windows are determined based on half-firing rate, when the firing rates were half of the peak firing rate).

The InRF and OutRF trials were further subgrouped on the basis of attentional selection. **Exo-InRF** trials are exo-conflict trials on which Poor-Bright target was selected and target was in the RF, whereas **Exo-OutRF** trials are exo-conflict trials on which Poor-Bright target was selected and target was out of the RF. Similarly, **Endo-InRF** trials are endo-conflict trials on which **Rich-Dim** target was selected and the target was in the RF, whereas **Endo-OutRF** trials are endo-conflict trials on which Rich-Dim target was selected and the target was out of the RF. The subgrouping of InRF and OutRF trials based on attentional selection yielded the following number of trials for each subgroup. The trials were pooled across neurons in three cell-type (VM, visual and movement neurons) groups. VM neurons: Exo-InRF=5459, Exo-OutRF=5379, Endo-InRF=16842, and Endo-OutRF=16470 trials. Visual neurons: Exo-InRF=2308, Exo-OutRF=2567, Endo-InRF=644 and Endo-OutRF=6740 trials. Movement neurons: Exo-InRF=2548, Exo-OutRF=2540, Endo-InRF=7388 and Endo-OutRF=7230 trials .

*Selection-time (ST) analysis:* We estimated the onset of selectivity in firing rates as the time after target onset when firing rates differed significantly for InRF and OutRF selection. We did this by first calculating the firing rates using a 15 ms smoothing window and then computing the difference in InRF and OutRF firing rates for each neuron. We tested the mean difference in firing rate for each group (**Fig 2E**) against a null hypothesis that there is no difference in firing rates using a permutation test. A null distribution of firing rate differences was generated by shuffling the InRF and OutRF firing rates across neurons in each group 1000 times. We detected ST as the first time-point when InRF firing rates were significantly greater than OutRF rates (p<0.01, permutation test). Since this procedure involves multiple comparisons, we corrected the p values by controlling for the false discovery rate.

*Spike-field coherence analysis:* We estimated spike-field coherence (SFC) as a function of frequency using multitaper spectral estimation (Mitra and Pesaran, 1999; Pesaran et al., 2002) with 10 Hz smoothing, and an estimation window spanning 200 ms before the target onset. The SFC was estimated between spiking and nearby LFP activity (within approx. 1.5 mm) to account for spiking activity bleeds into the LFP recording (**Supp. Fig. 23).** There was no spike amplitude for the broad-band recording on the LFP electrode when the activity was triggered on the spike times recorded on the other electrode.

The significance of SFC for each spike-field pair was tested against a null hypothesis that there was no SFC using a permutation test (1000 permutations, p<0.05). Null distribution for no SFC was generated by randomly permuting the order of trials for the spiking data compared to the LFP data. Raw coherence values were converted to z-scores by subtracting the mean and then dividing by the standard deviation of the null distribution. We applied cluster correction to identify the significant clusters of p-values while accounting for multiple comparisons (Maris et al., 2007). The significant cluster in beta (15-35 Hz) and gamma (40-70 Hz) frequency range was selected after performing a permutation test (1000 permutations, p<0.05). The coherent and not-coherent spike-field pairs in beta and gamma frequency ranges were identified based on the presence of a significant cluster in respective frequency bands. We identified 176 (M1=59, M2=117) coherent and 233 (M1=120, M2=113) not-coherent pairs in beta frequency range. A small number of spike-field pairs (10 out of 179 in M1 and 11 out of 230 in M2) were coherent in the gamma frequency range.

#### Beta-amplitude analysis

At each recording site, we tested whether beta bursts are specifically present in the 200ms prior to target onset against the null hypothesis of activity at other times during the trial using a permutation test. Across the population, beta bursts were reliably present in ∼96 % of recording sites in each animal (M1: 1108 out of 1152 sites; M2: 1299 out of 1344 sites; p<0.05, permutation test). We estimated amplitude of pre-target beta-burst for each trial at a single site, using multitaper spectral estimation (Mitra and Pesaran, 1999; Pesaran et al., 2002). We used 5 Hz smoothing, and an estimation window from 200 ms before target onset until target onset. The power values in beta (15-30 Hz) frequency range were converted to amplitude by taking square root. The logarithm transform of beta amplitude values were normalized with respect to mean across trials. We used these normalized beta-burst amplitude values for further analysis. Beta values varied trial-by-trial and observed a gaussian distribution at a given site (**Fig 3B**).

We examine the time-course of beta-burst related modulation in firing rates for exogenous and endogenous selection, by computing the beta values in six different 200 ms long time-windows (**Fig 6**). If otherwise mentioned beta related modulations were referred to beta values computed in 200 ms time-window before the target onset.

For a given site, we grouped the trials with the highest ∼33% and lowest ∼33% beta values to yield high-beta (HB) trials and low-beta (LB) trials. We calculated the SFC separately for HB and LB trials for coherent and not-coherent neurons.

#### Beta-bursts and attentional selection

We compared the firing responses of VM neurons on high-beta and low-beta trials for exogenous and endogenous selection. We further subgrouped the exo/endo InRF and OutRF trials based on beta values to yield the following number of trials for each subgroup. VM neurons -high-beta: Exo-InRF=1818, Exo-OutRF=1801, Endo-InRF=5599, Endo-OutRF=5458 trials, VM neurons-low-beta: Exo-InRF=1808, Exo-OutRF=1815, Endo-InRF=5524, Endo-OutRF=5513 trials. Similarly, for visual neurons we yielded, high-beta: Exo-InRF=792, Exo-OutRF=866, Endo-InRF=2166, Endo-OutRF=2222 trials and low-beta: Exo-InRF=772, Exo-OutRF=867, Endo-InRF=2110, Endo-OutRF=2306 trials. And for movement neurons we yielded, high-beta: Exo-InRF=852, Exo-OutRF=863, Endo-InRF=2459, Endo-OutRF=2345 trials and low-beta: Exo-InRF=849, Exo-OutRF=843, Endo-InRF=2468, Endo-OutRF=2445 trials.

#### Permutation test

We tested the difference in firing rates on HB and LB trials for each group in **Fig 4C, 5C** and **6C**. We computed the difference in firing rates between LB and HB trials for each neuron and tested the mean difference across neurons against a null hypothesis that there is no difference in firing rates using a permutation test. A null distribution of firing rates difference was generated by shuffling the HB and LB firing rates across neurons in each group 1000 times (p<0.01, permutation test). Since this procedure involves multiple comparisons, we corrected the p values by controlling for the false discovery rate.

We tested the difference in RTs after stratifying trials according to beta value for endogenous-conflict and exogeneous-conflict trials as shown in **Fig 7A**. We performed this test separately for each monkey. For each group of trials, we computed the test statistic given by the maximum difference in RT (range = max RT - min RT) after stratifying trials by beta value. We then tested the hypothesis that the difference in RT across beta values differed for the specific group of exogenous or endogenous trials using a permutation test. A null distribution of the test statistic was generated by shuffling the trial labels (endogenous-conflict and exogeneous-conflict) and then randomly assigning trials into groups of the same size as the original data set. We then computed the RT for trials stratified by beta values as for the original data set. RT for each group was standardized to be mean 1 before permuting by dividing by the mean RT in each group. Beta values for each group were standardized to be mean zero for each group before permuting by subtracting the mean beta value in each group. We performed this permutation 1000 times and compared the maximum difference in RT for each permutation with the test statistic (p<0.01, permutation test). We used an analogous procedure to test for a significant difference in RTs after stratifying trials according to beta value for luminance-only and reward-only trials, as shown in **Fig 7B**. Since this procedure was performed once per monkey and group, it was not necessary to control for multiple comparisons.

## Supplementary Information

**Supplementary Figure 1:**
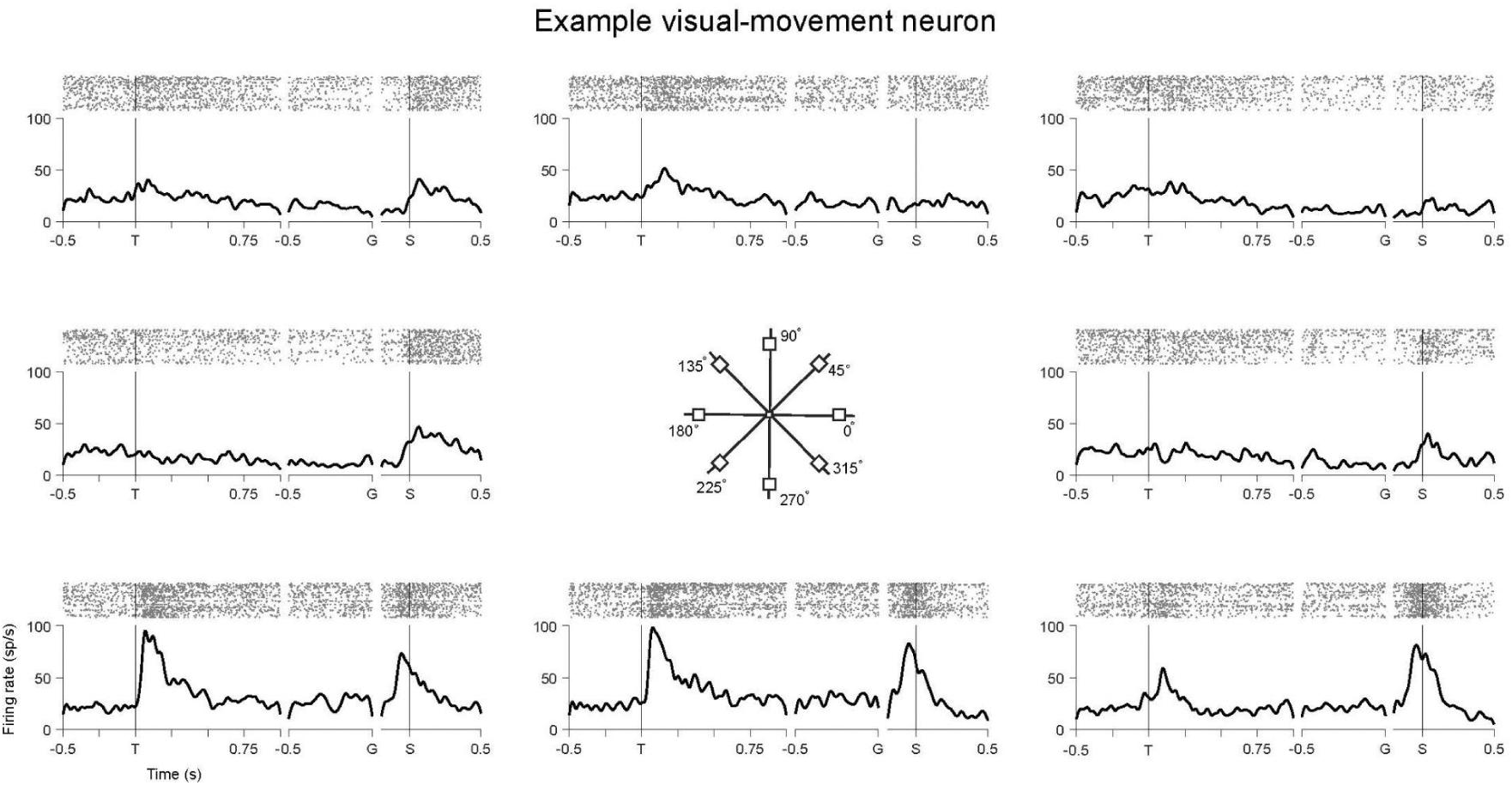
Example visual-movement neuron. Spike raters and peri-stimulus time histograms (PSTHs) of the example visual-movement neuron (same as **Fig 1E**) on ODR task trials. Each trial began with a visual target presented at the center of the screen for a variable 500-800 ms baseline period. After the baseline period, a red square appeared in one of the eight locations on a 10 deg circle in the visual periphery. After the variable 1000-1500 ms delay period, the central fixation target extinguished providing the Go signal to make a saccade to the target. T denotes the target onset, G denotes Go signal and S denotes Saccade Start.

**Supplementary Figure 2:**
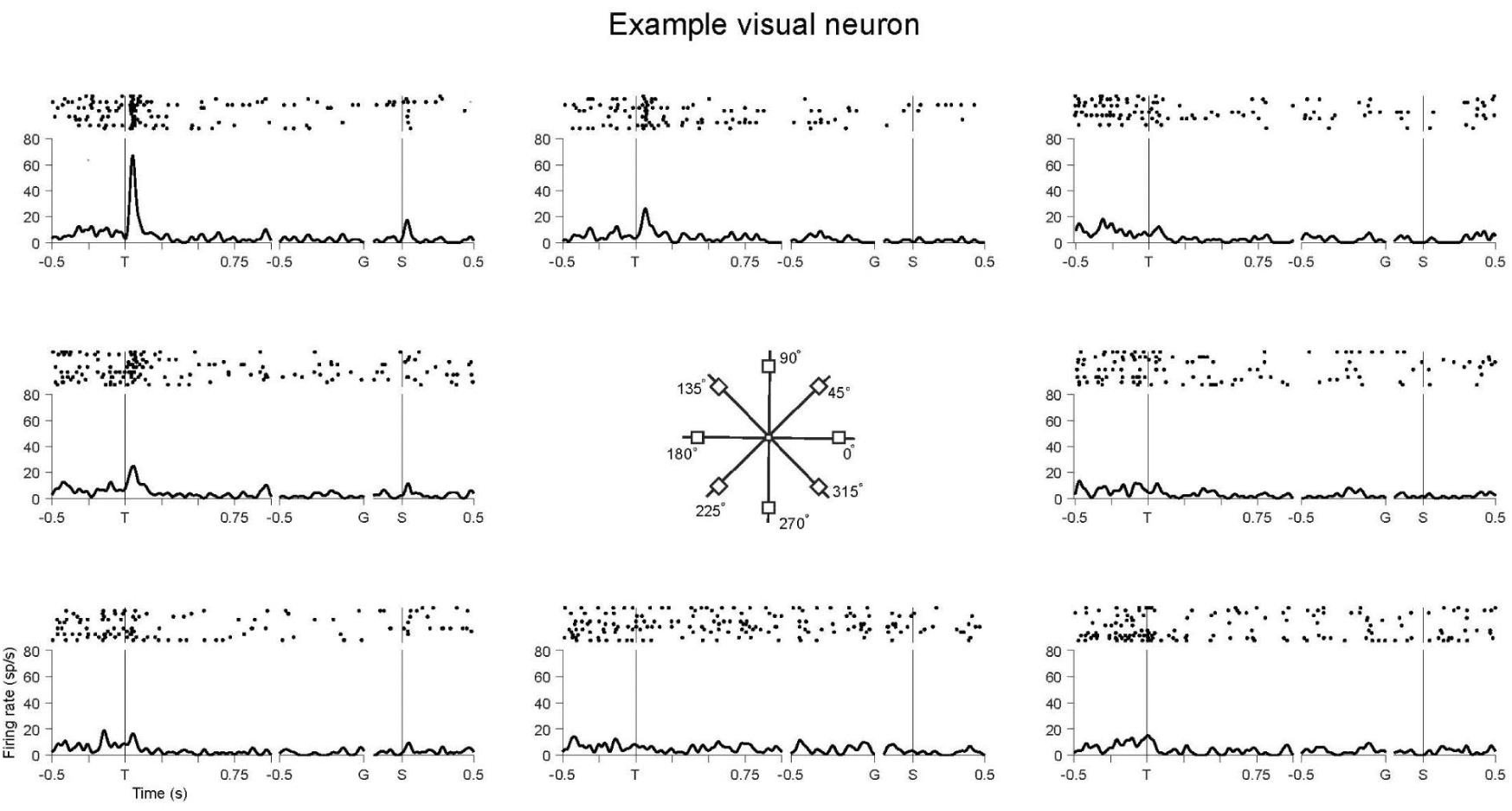
Example visual neuron. Spike raters and peri-stimulus time histograms (PSTHs) of the example visual neuron (same as **Fig 1E**) on ODR task trials. T denotes the target onset, G denotes Go signal and S denotes Saccade Start.

**Supplementary Figure 3:**
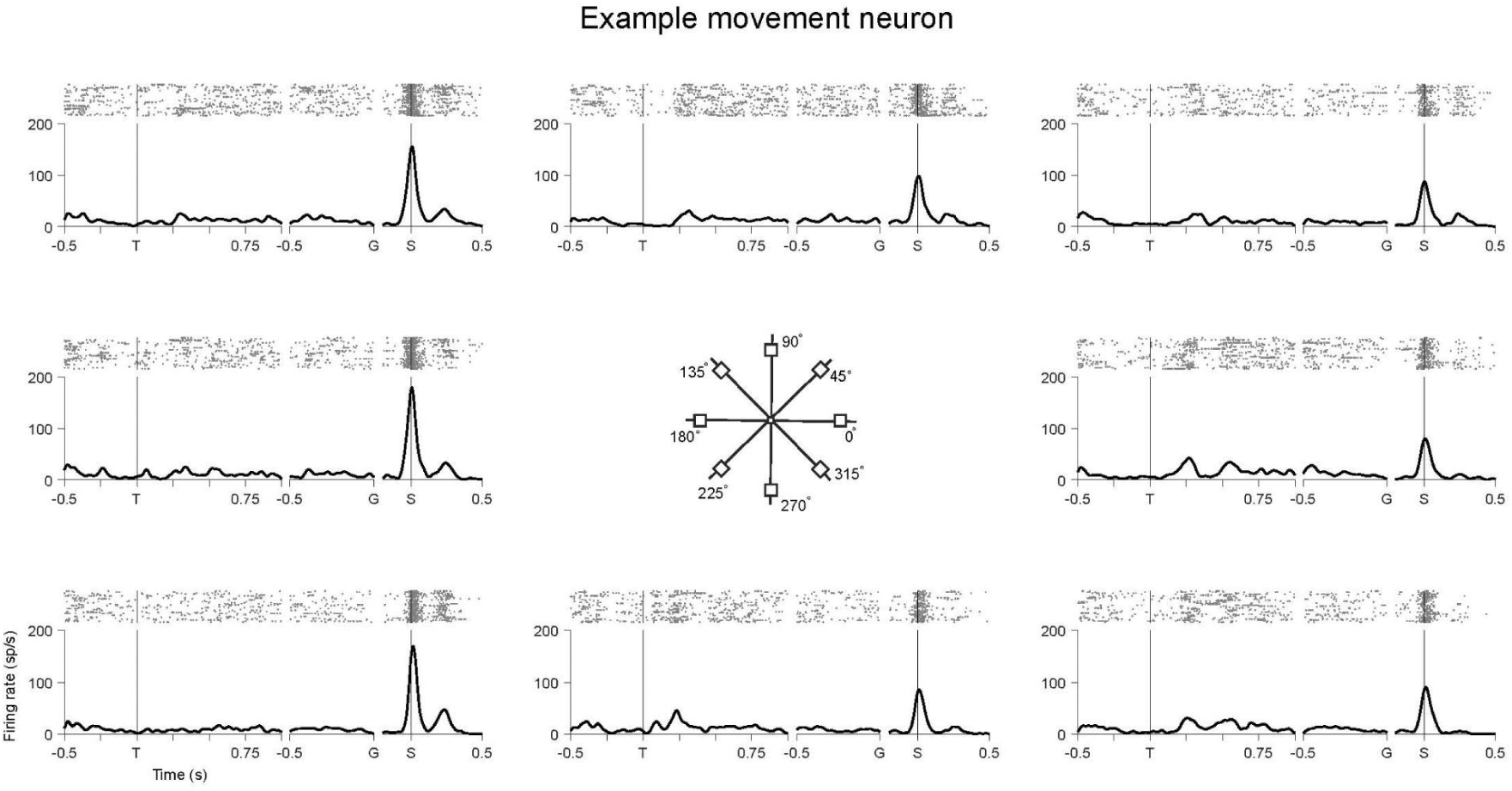
Example movement neuron. Spike raters and peri-stimulus time histograms (PSTHs) of the example visual neuron (same as **Fig 1E**) on ODR task trials. T denotes the target onset, G denotes Go signal and S denotes Saccade Start.

**Supplementary Figure 4:**
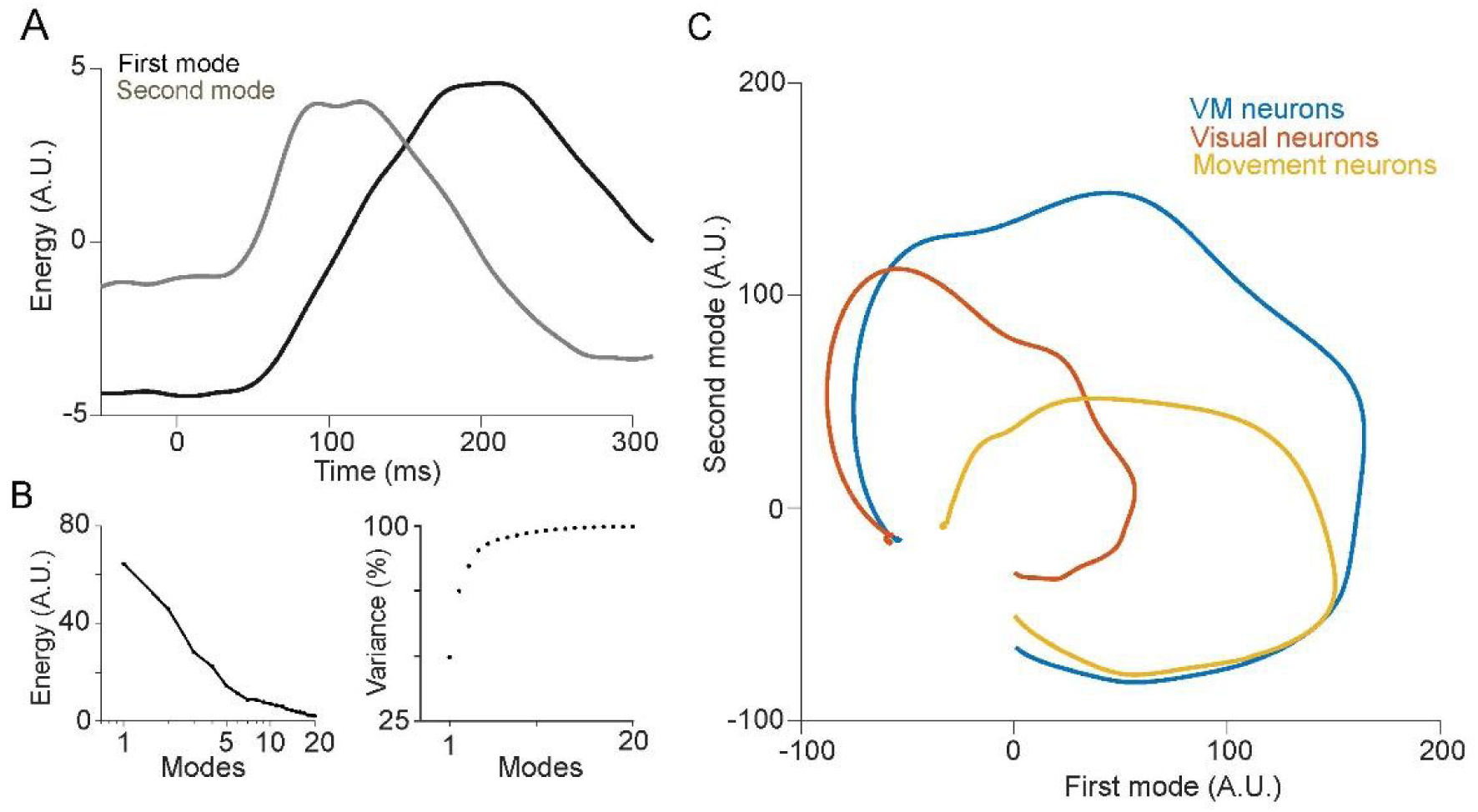
Neural population dynamics of LPFC neurons activity during LRS task (A) The first two modes obtained from principal component analysis (PCA) of LPFC neurons on luminance-only trials represent movement (first mode) and visual (second mode) activity**. (B)** Energy and variance explained by the first twenty modes of PCA. The first two modes explained ∼75 % of variability. **(C)** Projection of Visual-movement (blue), visual (red) and movement (yellow) neurons firing activity during independent set of LRS task trials on first two modes shown in A. VM neurons showed activity for both visual and movement modes whereas visual neurons showed activity for the second (visual) mode only. Similarly, movement neurons showed activity for the first (movement) mode only and not the visual mode.

**Supplementary Figure 5:**
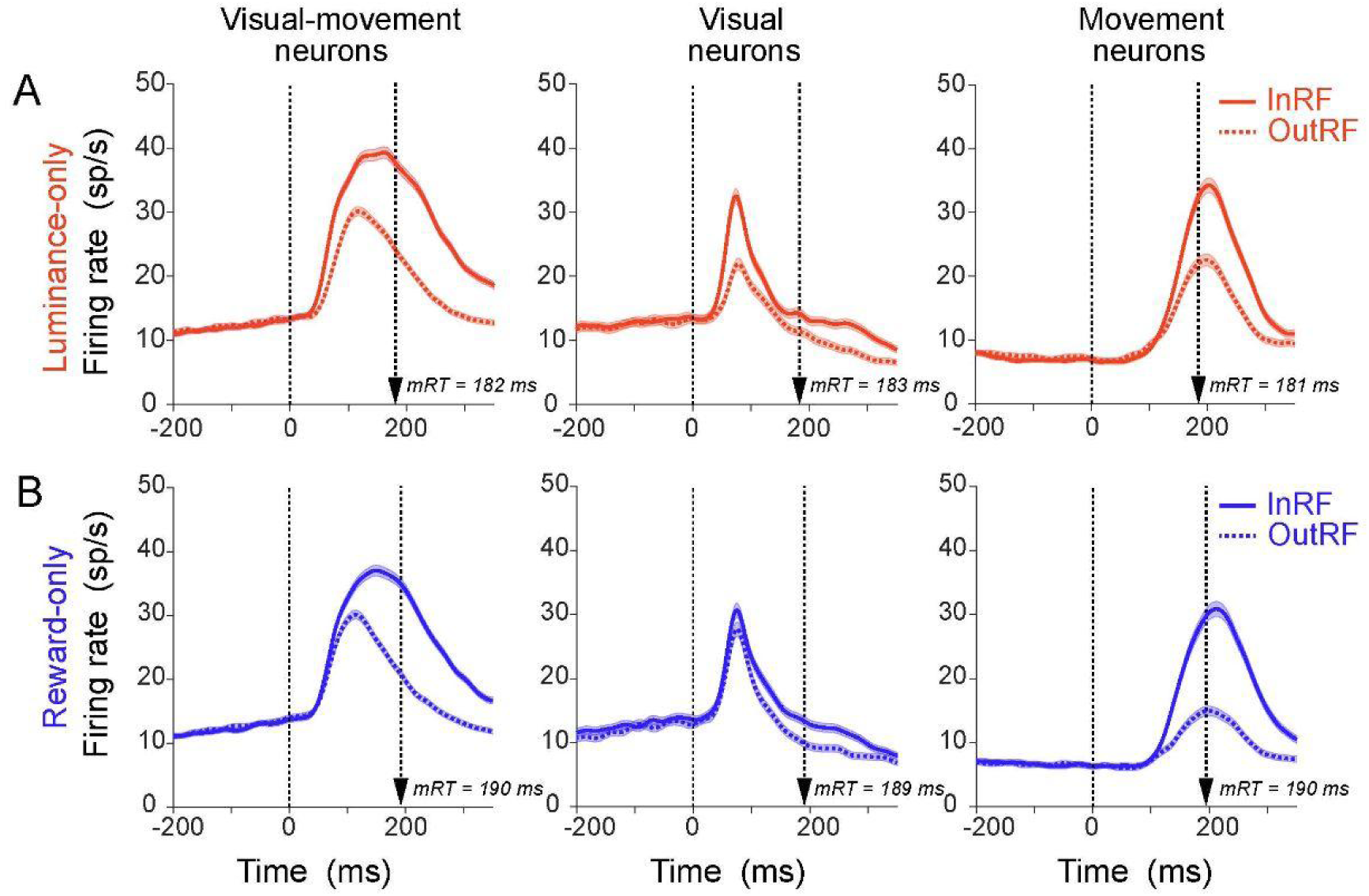
Population average firing rates of LPFC neurons on InRF and OutRF trials. **(A)** Population average firing rates of VM (N=139, left), visual (N=57, middle) and movement (N=65, right) neurons on luminance-only trials when **Bright** target was InRF (solid) and OutRF (dotted). The s.e.m of firing rates is shaded in lighter shades. **(B)** Same as A but for Reward-only trials when Rich target was InRF (solid) and OutRF (dotted). Dotted lines denote target onset.

**Supplementary Figure 6:**
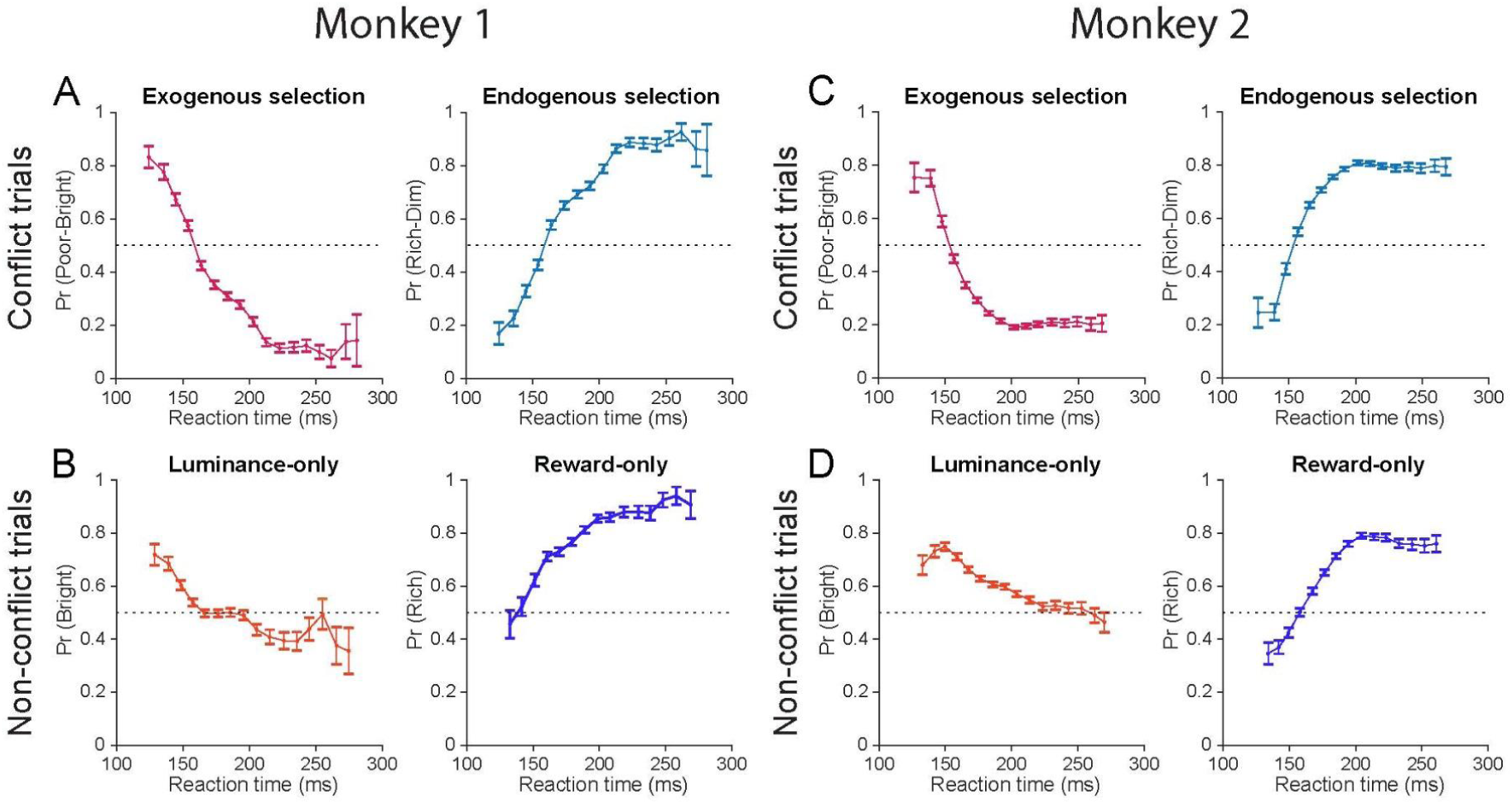
Choice behavior on conflict and non-conflict trials (A) Target choice probabilities on LRS conflict trials for Monkey 1. Poor-Bright target choice probability reflected exogenous selection and Rich-Dim target choice probability reflected endogenous selection. Note that Pr (Poor-Bright) is 1 - Pr (Rich-Dim) **(B)** Target choice probabilities on luminance-only (left panel) and reward only (right panel) non-conflict trials. Bright target choice probability predominantly reflected exogenous selection on shorter RT trials. Rich target choice probability predominantly reflected endogenous selection on longer RT trials **(C,D)** Same as A,B but for Monkey 2.

**Supplementary Figure 7:**
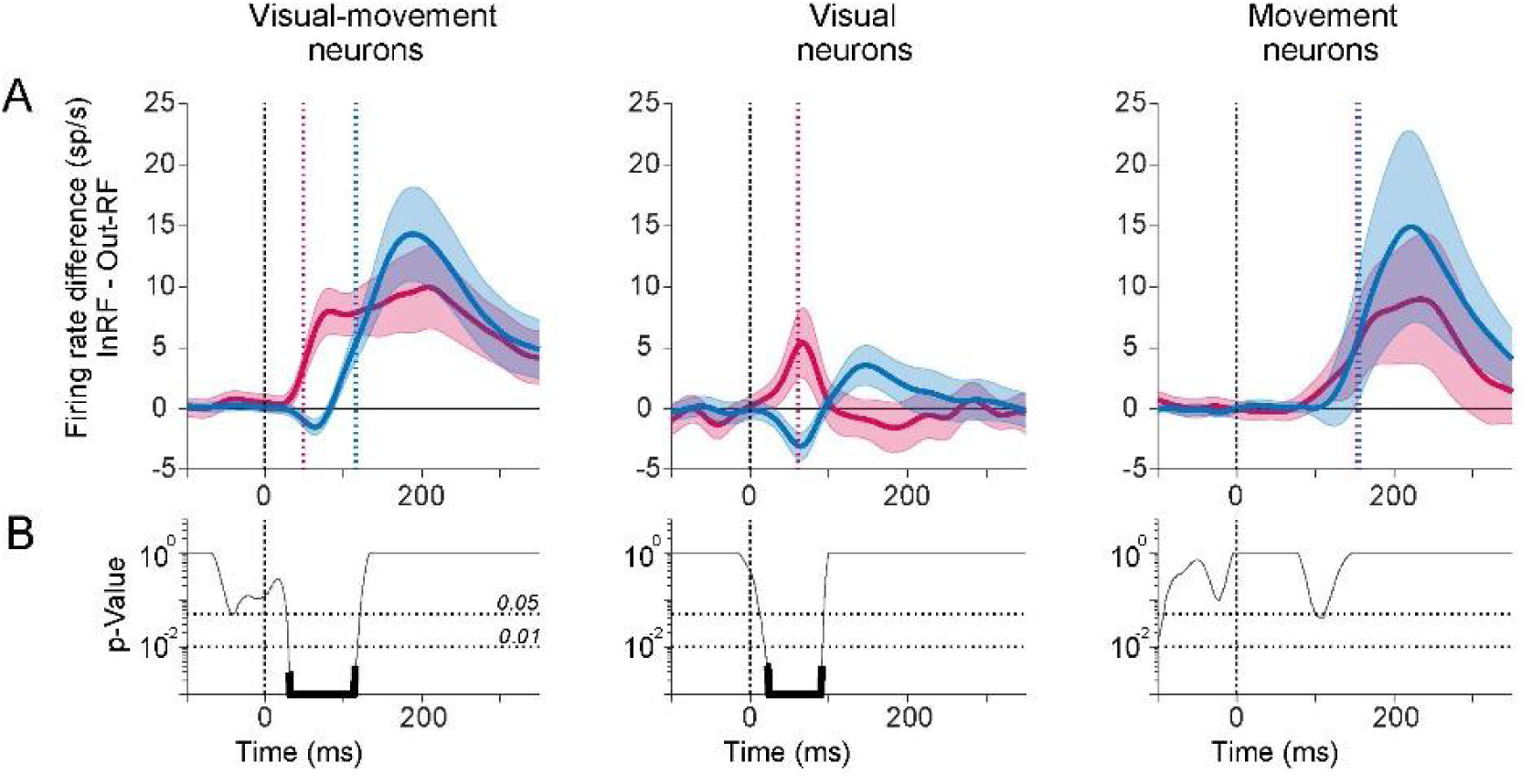
(A) Average firing rates differences between InRF and OutRF trials across VM neurons (N=139), visual neurons (N=57) and movement neurons (N=65). Exogenous selection is shown in red and endogenous selection in blue. The s.e.m of firing rate differences are shown in lighter shades. Black dotted lines denote target onset. Red dotted lines denote exogenous selection time and blue dotted lines denote endogenous selection time. **(B)** Permutation test p-values against a null hypothesis that differences in firing rates for exogenous and endogenous selection are similar are shown in black. The null distribution for the permutation test was generated by shuffling the endogenous and exogenous labels and then randomly assigning the firing rate differences into endogenous and exogenous groups of the same size as the original dataset. The endogenous firing rates were subtracted from the exogenous firing rate for each neuron session and an average across population was computed to yield firing rates differences. The permutation was performed 1000 times and compared with the test statistic to yield p-values. FDR corrected p-values for alpha = 0.01 is shown in bold.

**Supplementary Figure 8:**
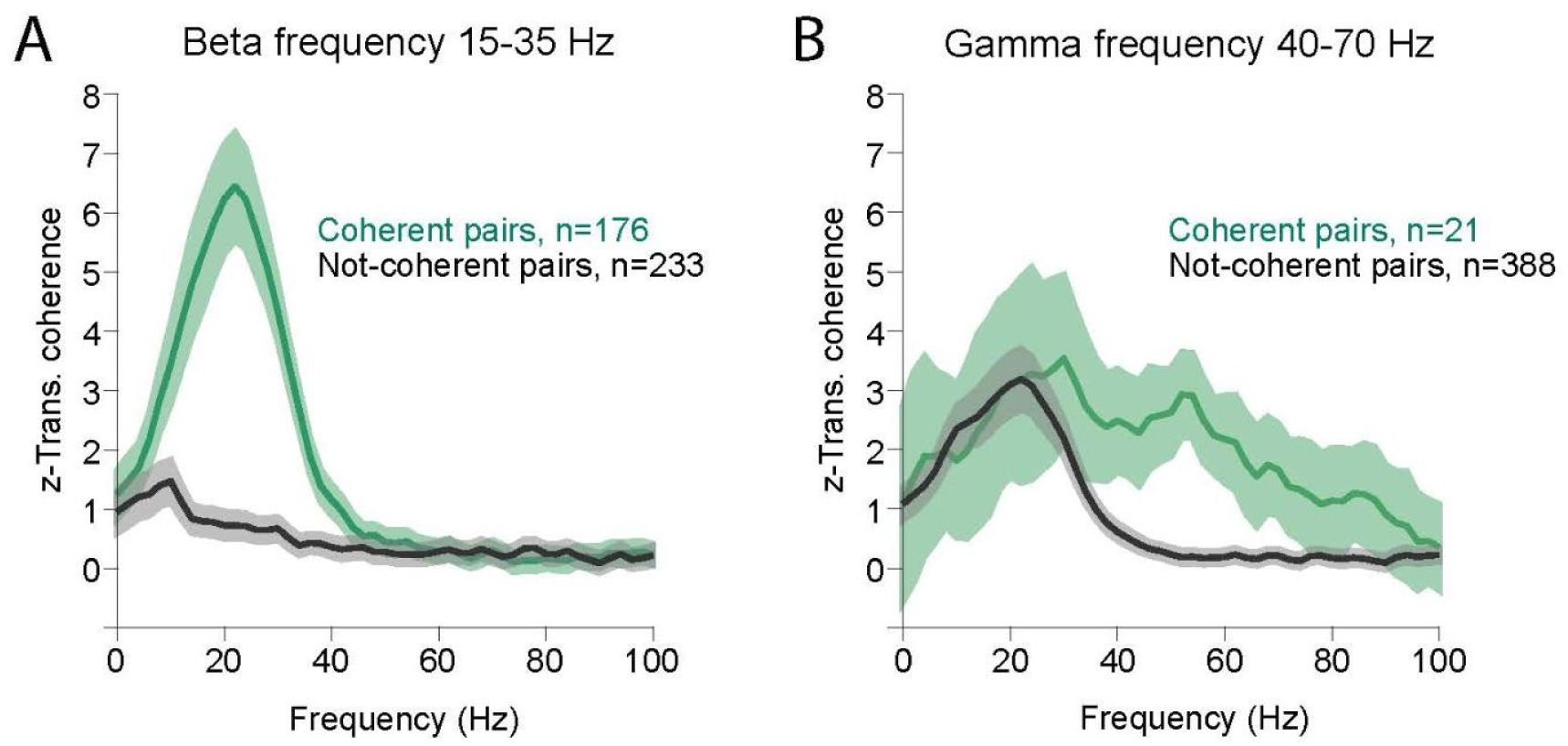
**(A)** Population average spike field coherence (SFC) of coherent and not-coherent pairs in beta frequency range (15-35 Hz). (**B)** Population SFC of coherent and not-coherent pairs in gamma frequency range (40-70 Hz). A small number (< 5%) of LPFC neurons fired coherently in the gamma range. The s.e.m. of SFC is shown in lighter shades.

**Supplementary Figure 9:**
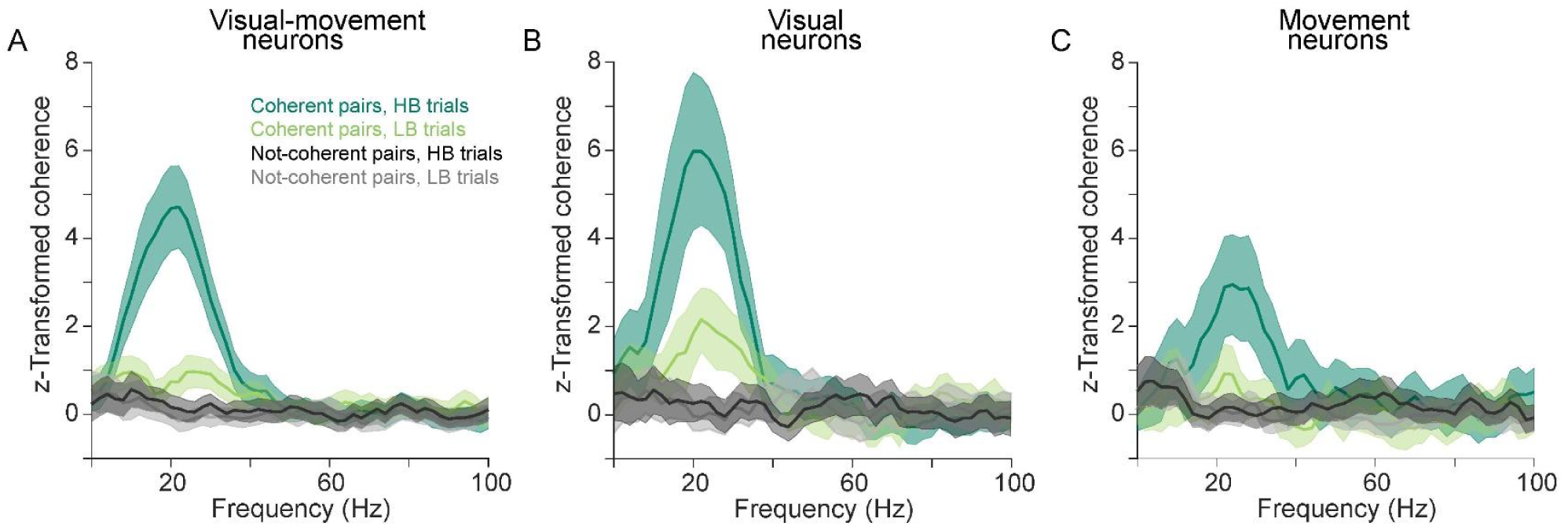
**(A)** Population average spike field coherence (SFC) of coherent (N=62) and not-coherent (N=77) pairs for visual-movement neurons in beta frequency range (15-35 Hz). The s.e.m of SFC is shown in lighter shades (**B)** Same as A, but for visual coherent (N=24) and not-coherent (N=33) neurons (C) Same as A, but for movement coherent (N=17) and not-coherent (N=48) neurons. The s.e.m. of SFC is shown in lighter shades.

**Supplementary Figure 10:**
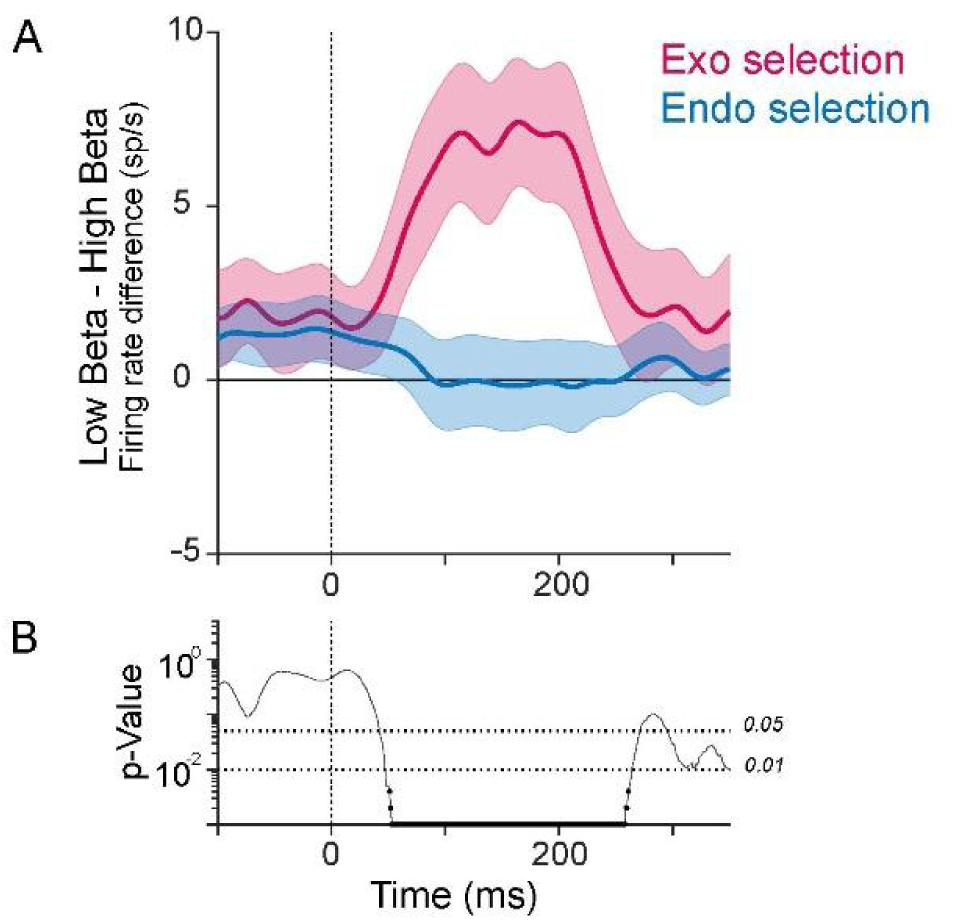
(A) Visual-movement neurons population average firing rate differences between low-beta and high-beta trials. Mean +/- s.e.m. are shown for exogenous (red) and endogenous (blue) selection InRF LRS conflict trials. **(B)** Exogenous and endogenous conditions are compared using a permutation against the null hypothesis that there is no difference between the two conditions. The null distribution is generated by shuffling the endogenous and exogenous labels and then randomly assigning the firing rate differences into endogenous and exogenous groups of the same size as the original dataset. The endogenous firing rates were subtracted from the exogenous firing rate for each neuron session and an average across population was computed to yield firing rates differences. The permutation was performed 1000 times and compared with the test statistic to yield p-values. FDR corrected p-values for alpha = 0.01 is shown in bold. Dotted lines denote target onset.

**Supplementary Figure 11:**
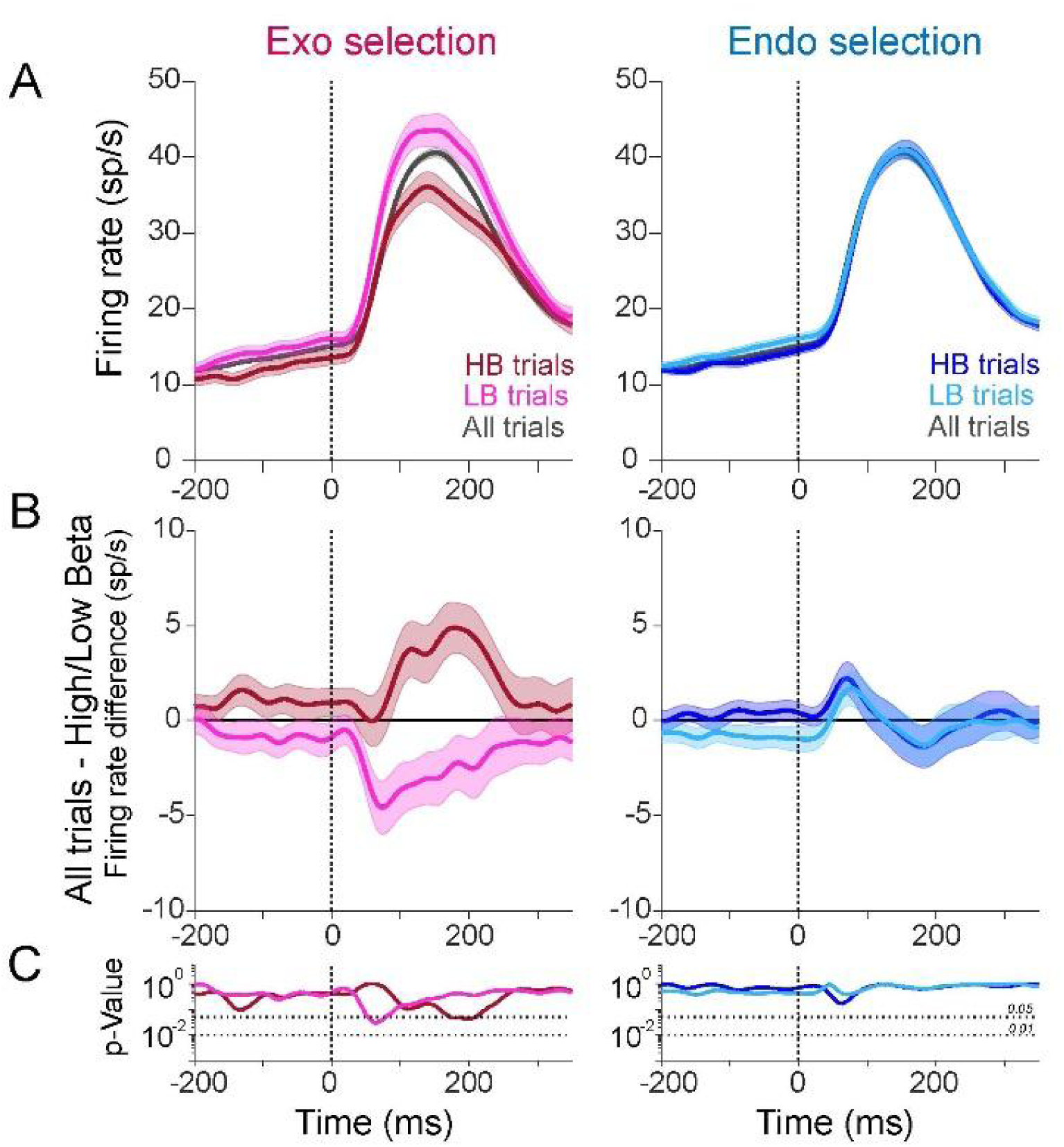
**(A)** Population average firing rates of VM neurons for exogenous (left panel) and endogenous (right panel) selection on InRF LRS conflict trials (All trials, black). High-beta trials (HB, darker traces) represent InRF trials when pre-target beta-burst is high and the low-beta trials (LB, lighter traces) pre-target beta-burst is low. **(B)** Difference in firing rates for all trials and HB trials (darker traces) and all traces and LB (lighter traces). Mean +/- s.e.m. are shown for exogenous (red) and endogenous (blue) selection. **(C)** Permutation test p-values under a null hypothesis that there is no difference in firing rates for all and high/low beta trials. Dotted lines denote target onset.

**Supplementary Figure 12:**
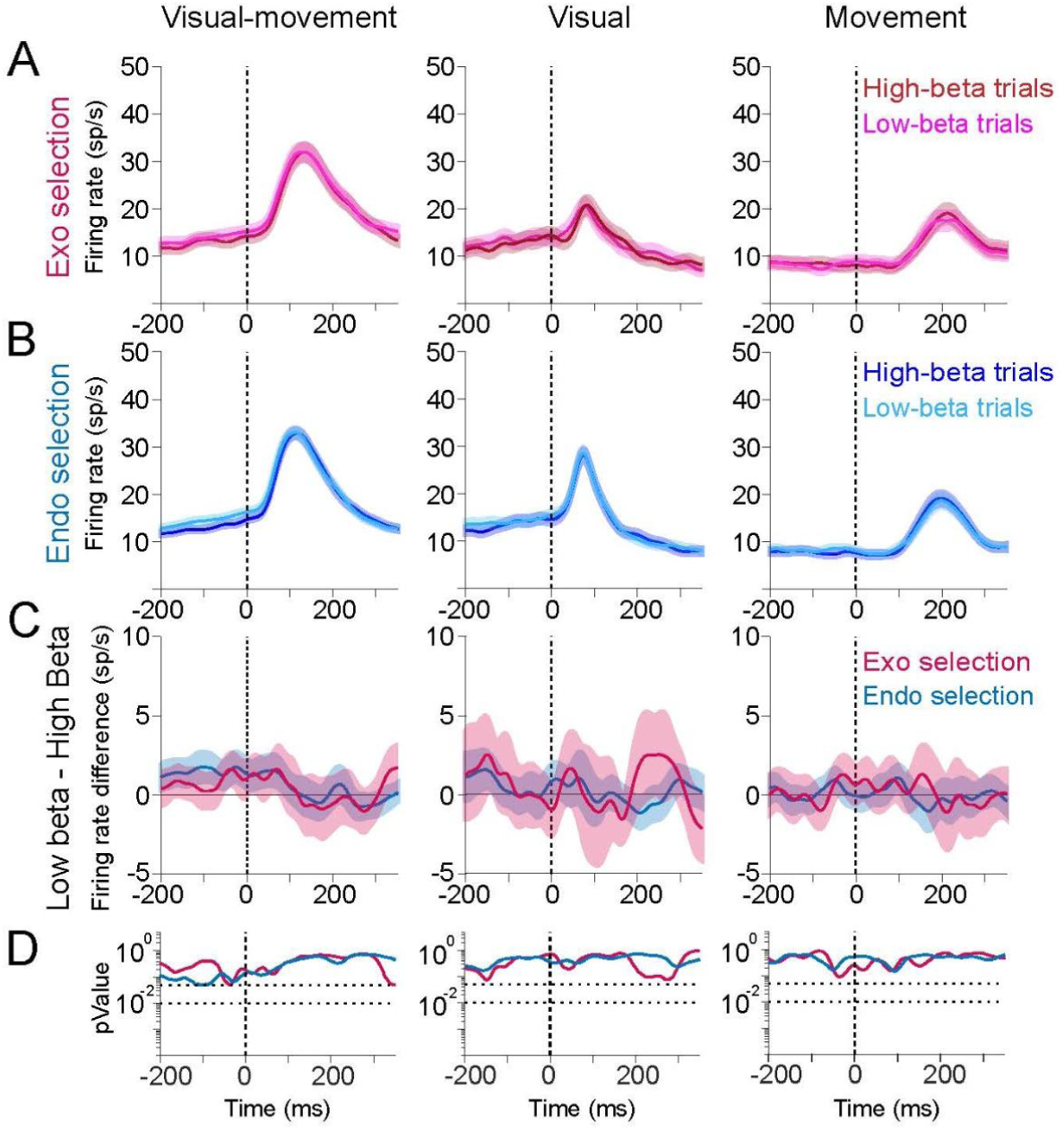
Beta-bursts do not modulate neuronal firing when attentional selection is outside the response field. **(A)** Population average firing rates of the VM, visual and movement neurons for exogenous selection when pre-target beta-burst is high (HB trials, darker traces) and low (LB trials, lighter traces). Mean +/- s.e.m. are shown for OutRF trials when selection was outside the RF of the units. Dotted lines denote target onset. **(B)** Same as A but for endogenous selection. **(C)** Difference in firing rates for pre-target low and high beta-bursts. Mean +/- s.e.m. are shown for exogenous (red) and endogenous (blue) selection. **(D)** Permutation test p-values under a null hypothesis that there is no difference in firing rates for high-beta and low-beta trials for exogenous (red) and endogenous (blue) selection.

**Supplementary Figure 13:**
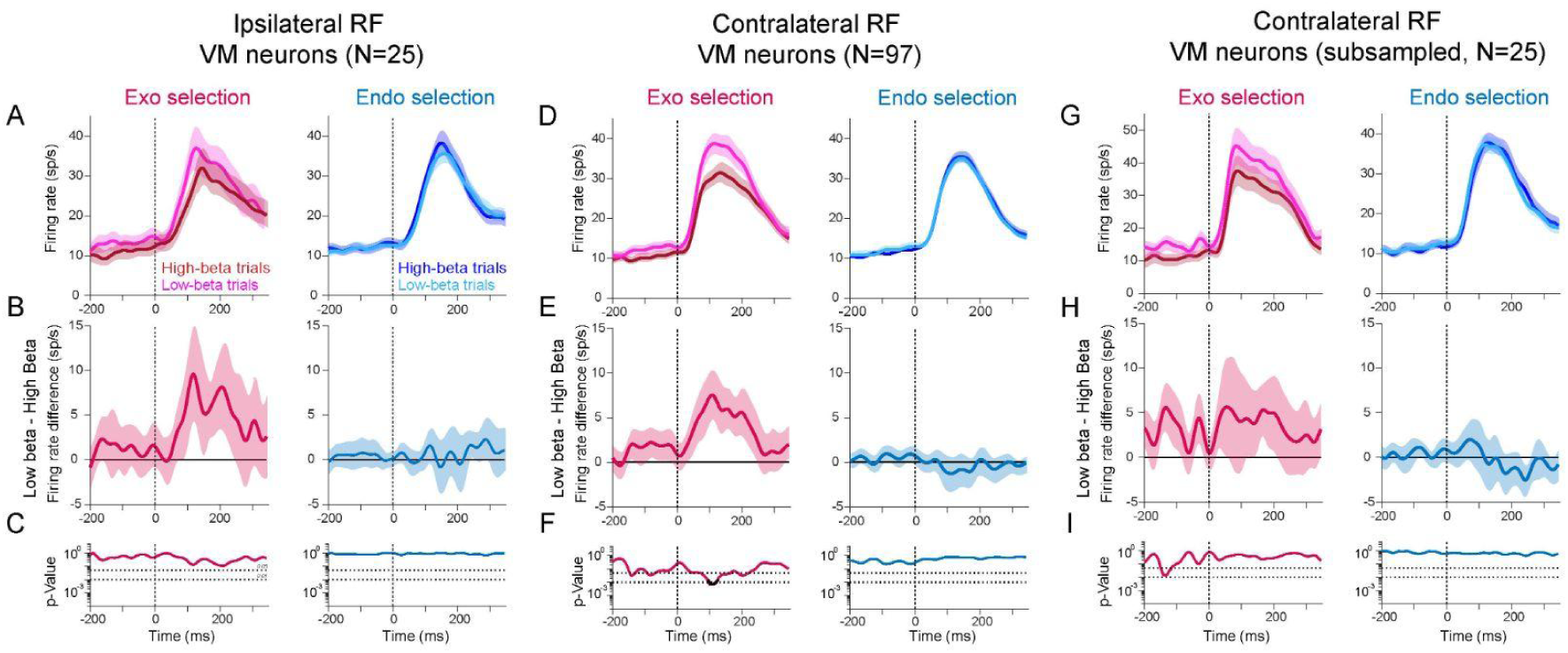
**(A)** Population average firing rates of VM neurons that have ipsilateral response fields (N=25) for exogenous (left panel) and endogenous (right panel) selection when pre-target beta is high (darker traces) and low (lighter traces). **(B)** Difference in firing rates for pre-target low and high beta-bursts. Mean +/- s.e.m. are shown for exogenous (red) and endogenous (blue) selection InRF LRS conflict trials. **(C,F,I)** Permutation test p-values under a null hypothesis that there is no difference in firing rates for high-beta and low-beta trials for exogenous (red) and endogenous (blue) selection. FDR corrected p-values for alpha = 0.01 is shown in bold. Dotted lines denote target onset. **(D,G)** Same as A but for VM neurons that have contralateral response fields (N=97) for total population and randomly selected N=25 contralateral-RF VM neurons. **(E,H)** Same as B but for contralateral-RF VM neurons and subsampled contralateral.

**Supplementary Figure 14:**
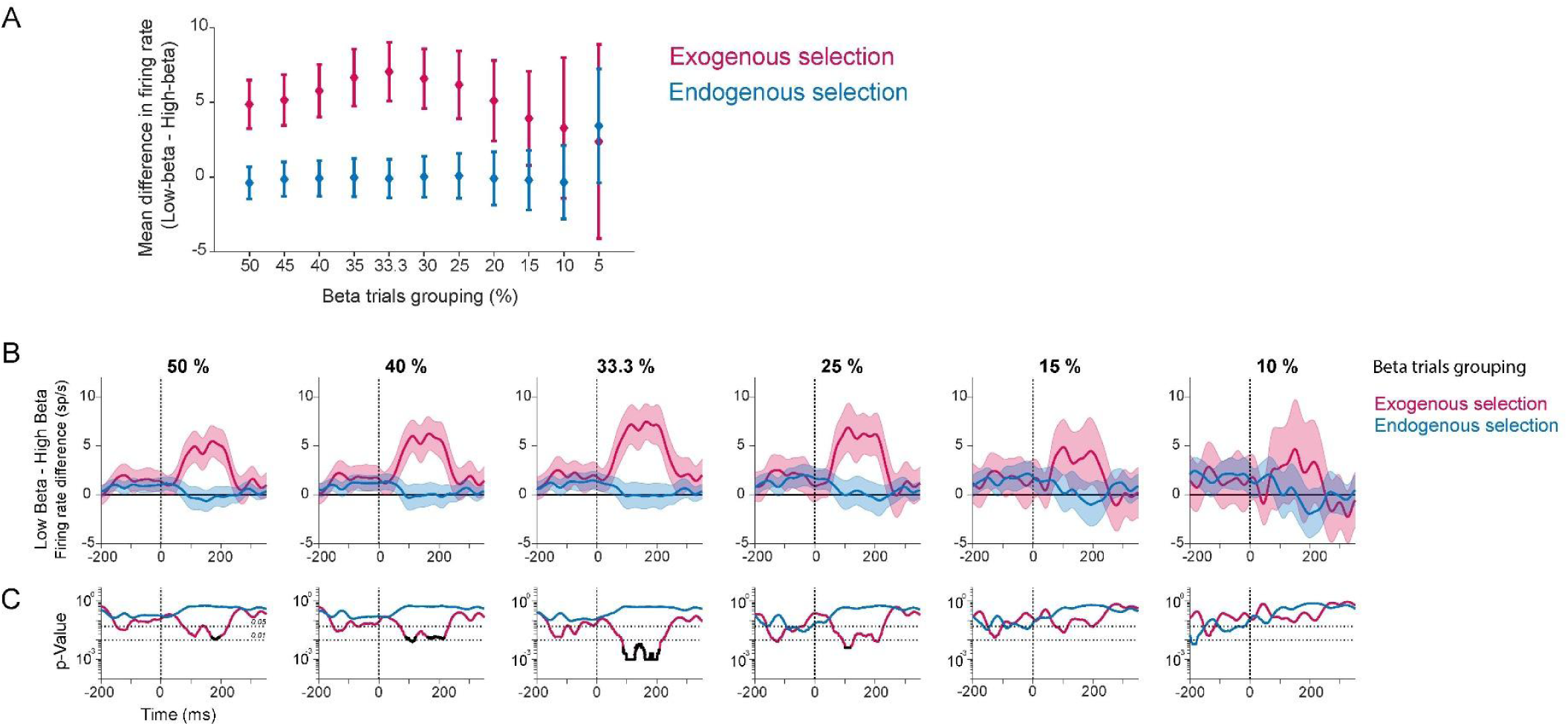
Beta-bursts effect on attentional selection for different beta trials grouping percentiles (A) Average difference in VM neurons firing rates for low and high beta trials for different beta trials grouping percentiles. Firing rate differences were averaged between 100 to 200 ms epoch for exogenous (red) and endogenous (blue) selection. **(B)** Difference in firing rates for pre-target low and high beta-bursts. Mean +/- s.e.m. are shown for exogenous (red) and endogenous (blue) selection InRF LRS conflict trials. **(C)** Permutation test p-values under a null hypothesis that there is no difference in firing rates for high-beta and low-beta trials for exogenous (red) and endogenous (blue) selection. FDR corrected p-values for alpha = 0.01 is shown in bold.

**Supplementary Figure 15:**
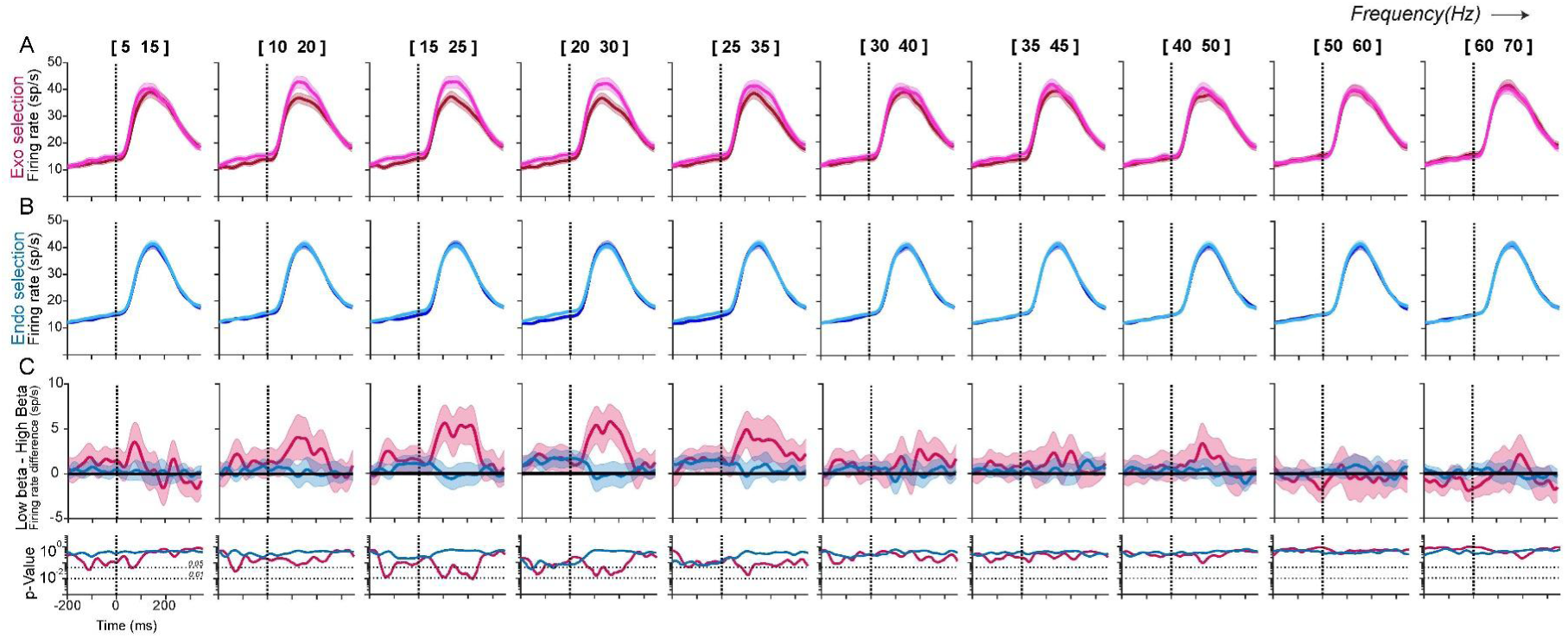
Pre-target LFP activity effect on attentional selection for different frequency ranges (A) Population average firing rates of the VM neurons for exogenous selection when pre-target beta-burst is high (HB trials, darker traces) and low (LB trials, lighter traces). Mean +/- s.e.m. are shown for InRF trials when exogenous selection was inside the RF. Dotted lines denote target onset. **(B)** Same as A but for endogenous selection. **(C)** Difference in firing rates for pre-target low and high beta-bursts. Mean +/- s.e.m. are shown for exogenous (red) and endogenous (blue) selection. **(D)** Permutation test p-values under a null hypothesis that there is no difference in firing rates for high-beta and low-beta trials for exogenous (red) and endogenous (blue) selection.

**Supplementary Figure 16:**
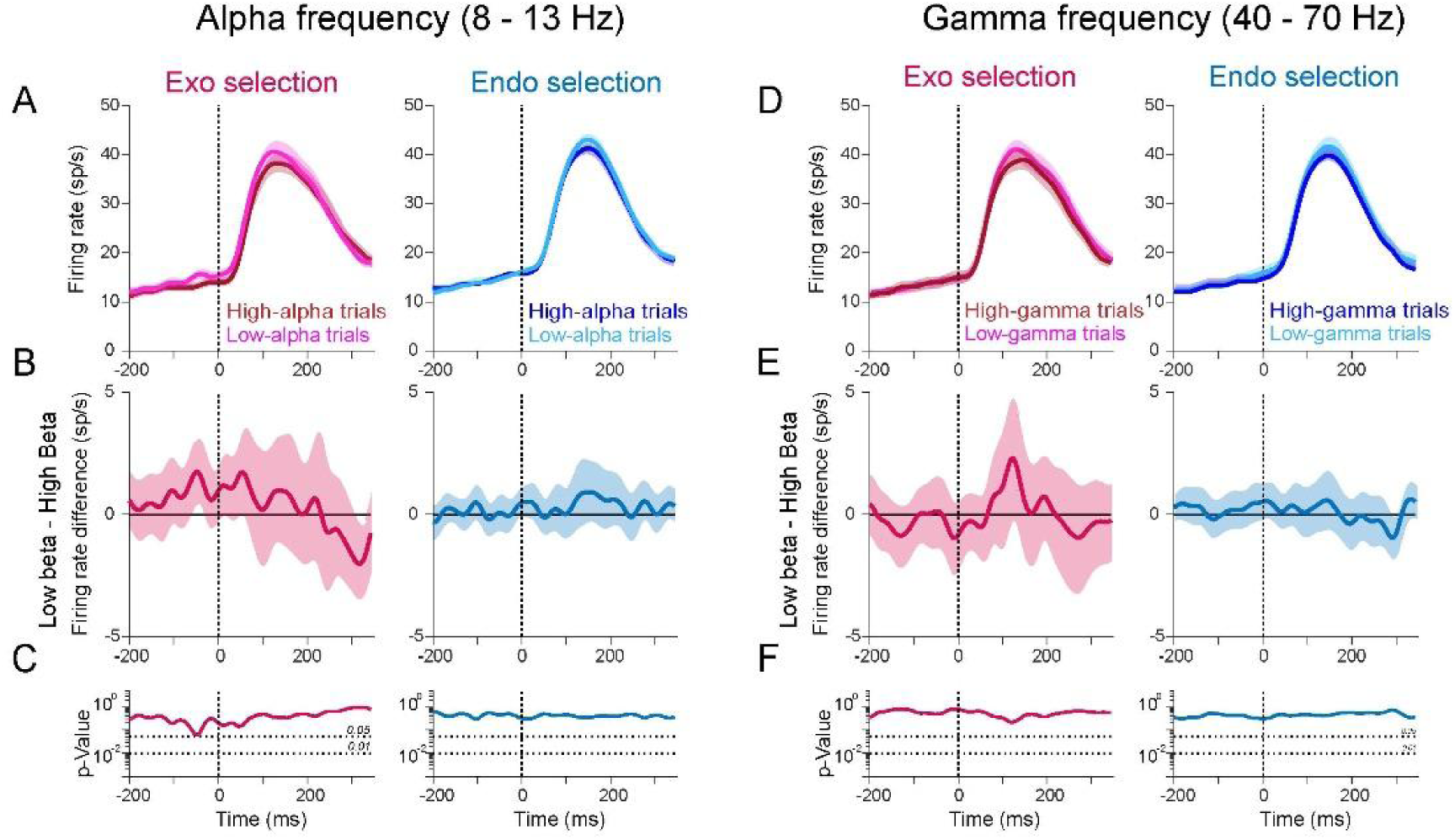
Pre-target alpha and gamma activity effect on attentional selection (A) Population average firing rates of the VM neurons for exogenous selection when pre-target alpha activity is high (darker traces) and low (lighter traces). Trials are grouped into the highest ∼33% and lowest ∼33% pre-target alpha range activity. Mean +/- s.e.m. are shown for InRF trials when exogenous (left panel) and endogenous (right) selection was inside the RF. Dotted lines denote target onset. **(B)** Difference in firing rates for pre-target low and high alpha activity. Mean +/- s.e.m. are shown for exogenous (red) and endogenous (blue) selection. **(C)** Permutation test p-values under a null hypothesis that there is no difference in firing rates for high-alpha and low-alpha trials for exogenous (red) and endogenous (blue) selection. **(D, E, F)** Same as A,B and C but for gamma frequency range.

**Supplementary Figure 17:**
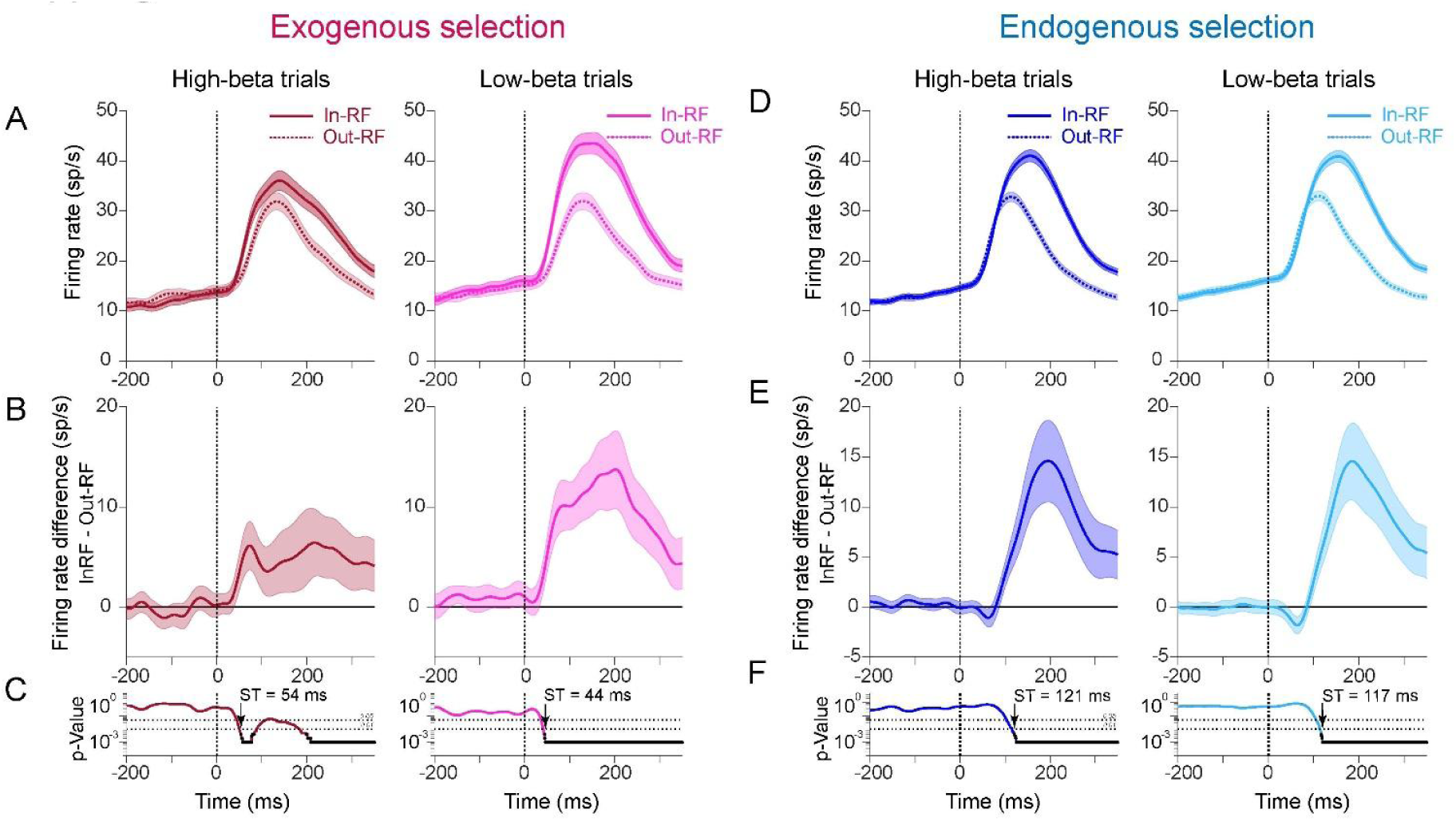
Selection time analysis for high and low beta trials. **(A)** Population average VM neurons firing rates for exogenous selection when pre-target beta is high (left panel) and low (right panel) trials. Mean +/- s.e.m. are shown for InRF trials (solid traces) and OutRF (dotted traces) trials. **(B)** Difference in firing rates for exogenous selection into and out of the RF for high and low beta groups. Mean +/- s.e.m. are shown **(C)** Permutation test p-values against a null hypothesis that there is no difference in InRF and OutRF firing rates. FDR corrected p-values for alpha = 0.01 are shown in black. Arrow represents the selection time (ST) when first time separation becomes significant. Exo-HB ST=54 ms, Exo-LB ST=44 ms. **(D, E, F)** Same as A, B and C but for endogenous selection. Endo-HB ST=121 ms, Endo-LB ST=117 ms.

**Supplementary Figure 18:**
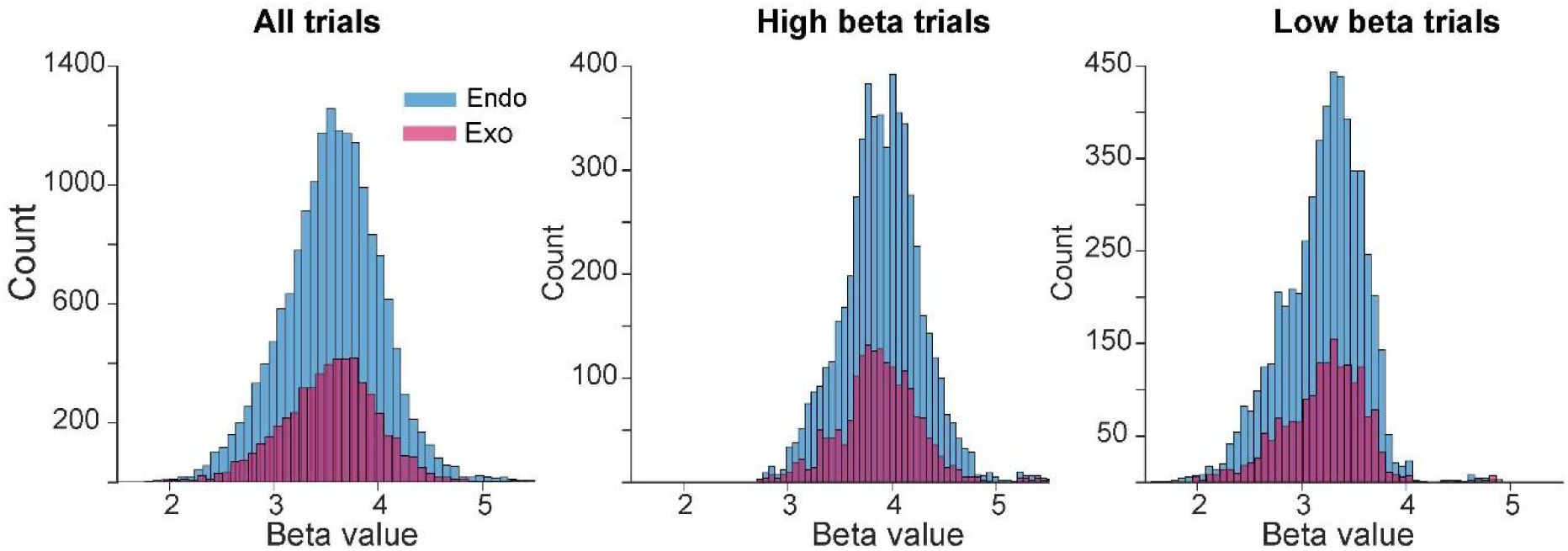
(A) Pre-target beta-burst values distribution for LRS conflict exogenous (red) and endogenous (blue) selection into the RF. **(B,C)** Same as A but for trials when pre-target beta-bursts amplitude is high and low. Pre-target beta-burst value for each trial at a single site was computed using multitaper spectral estimation (Mitra and Pesaran, 1999; Pesaran et al., 2002). We used a 5 Hz smoothing and an estimation window of 200 ms (before target onset until target onset). Beta values shown here are the logarithmic transform of average power values in beta (15-30 Hz) frequency range.

**Supplementary Figure 19:**
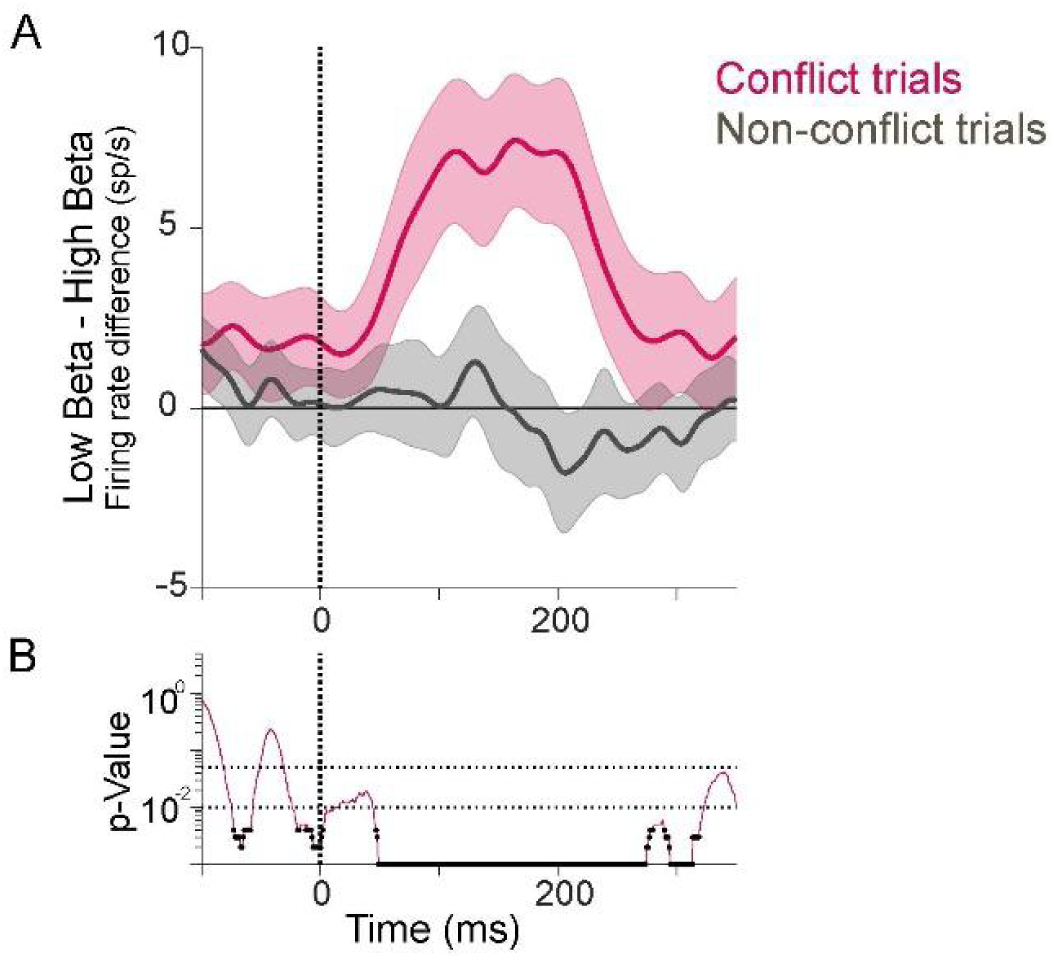
**(A)** Visual-movement neurons population average firing rate differences between low-beta and high-beta trials. Mean +/- s.e.m. are shown for exogenous-conflict trials (red) and luminance-only non-conflict trials (gray). **(B)** Conflict and non-conflict conditions are compared using a permutation against the null hypothesis that there is no difference between the two conditions. The null distribution is generated by shuffling the conflict and non-conflict labels and then randomly assigning the firing rate differences into conflict and non-conflict groups of the same size as the original dataset. The difference between two conditions is obtained for each neuron session and an average across population was computed to yield firing rates differences. The permutation was performed 1000 times and compared with the test statistic to yield p-values. FDR corrected p-values for alpha = 0.01 is shown in bold. Dotted lines denote target onset.

**Supplementary Figure 20:**
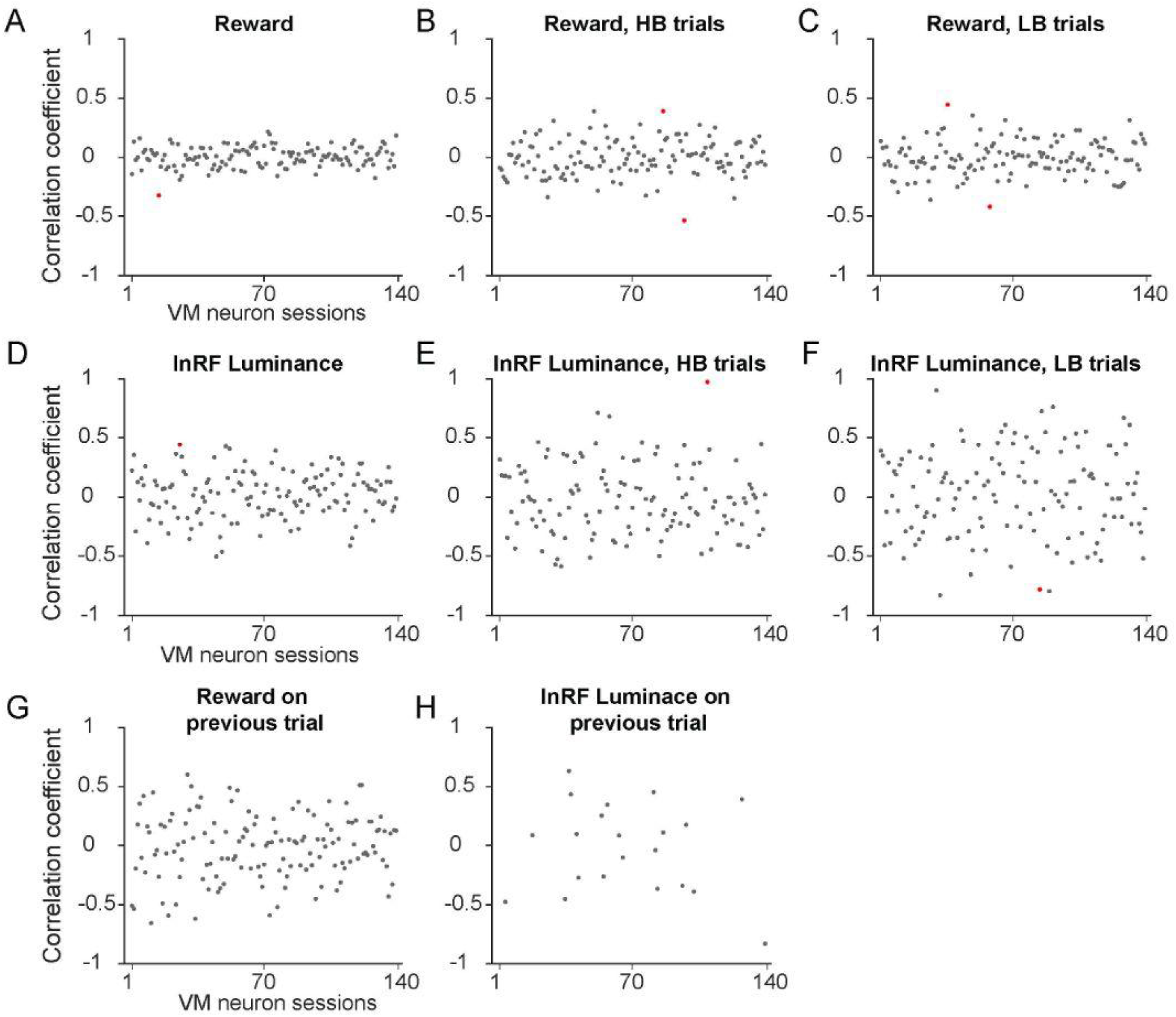
Pearson correlation coefficient between beta-burst amplitude and **(A)** reward value of the selected target on InRF trials **(B)** reward value of the selected target on high-beta InRF trials **(C)** reward value of the selected target on low-beta InRF trials **(D)** luminance value of the InRF stimulus **(E)** luminance value of the InRF stimulus on high-beta trials **(F)** luminance value of the InRF target on low-beta trials **(G)** reward value on the previous trial and **(H)** InRF luminance on previous trial for each of the VM neurons session (N=139). Since these analysis are performed for each session, sessions with at least 5 trials per condition (*i.e.* HB trials, LB trials or all trials) were analyzed. Across VM sessions the correlation coefficients ranged between negative to positive values and majority were not significant. The significant sessions after FDR correction are shown in red.

**Supplementary Figure 21:**
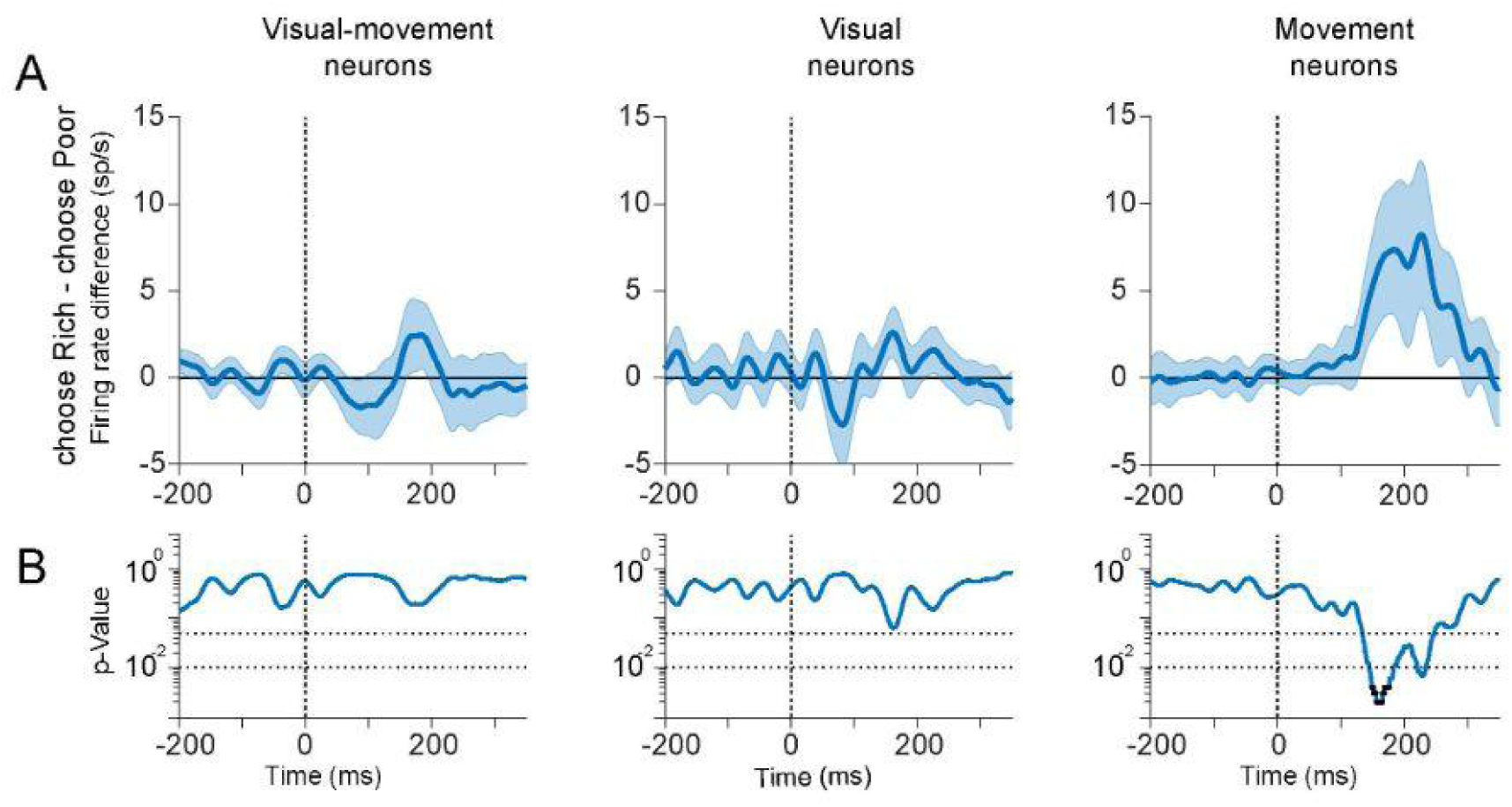
**(A)** Population average firing rate differences between Rich target and Poor target selection inside the RF. Mean +/- s.e.m. are shown for reward-only non-conflict InRF trials. **(B)** Permutation test p-values against a null hypothesis that there is no difference in firing rates between Rich and Poor target selection. FDR corrected p-values for alpha = 0.01 are shown in black.

**Supplementary Figure 22:**
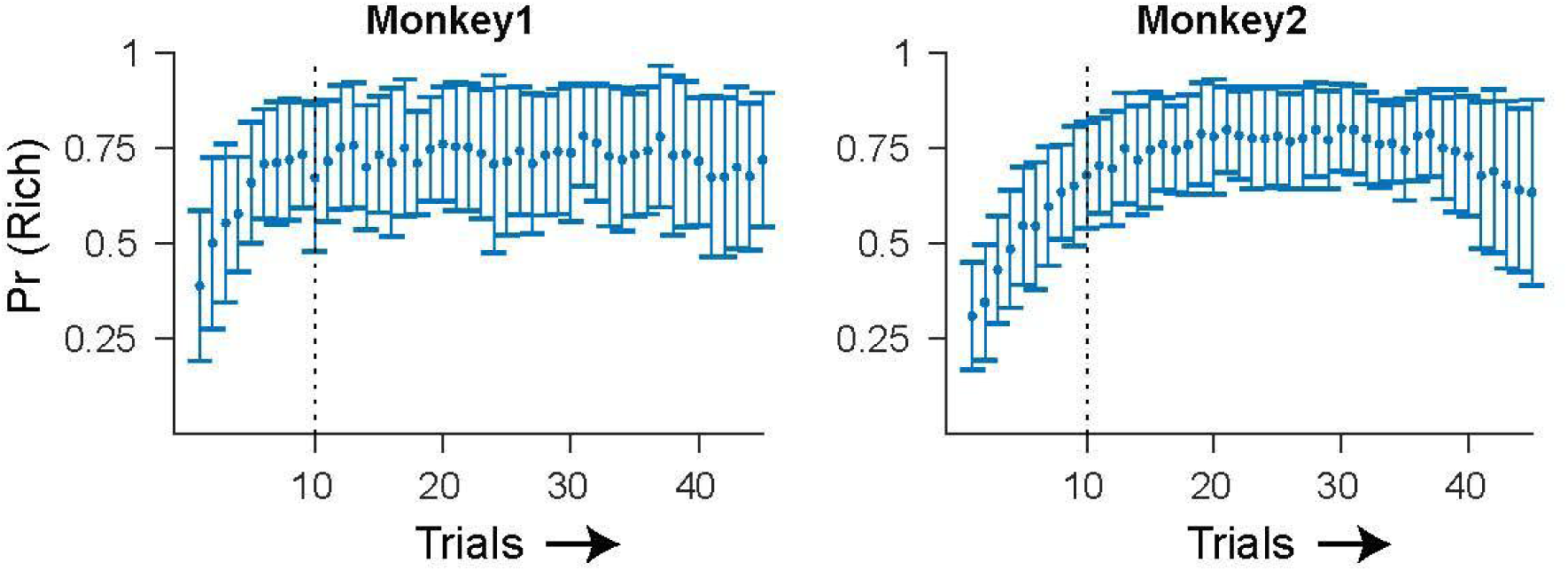
Choice behavior trial by trial around each reward block transition. Mean +/- s.e.m of Rich target choice probability across reward blocks.

**Supplementary Figure 23:**
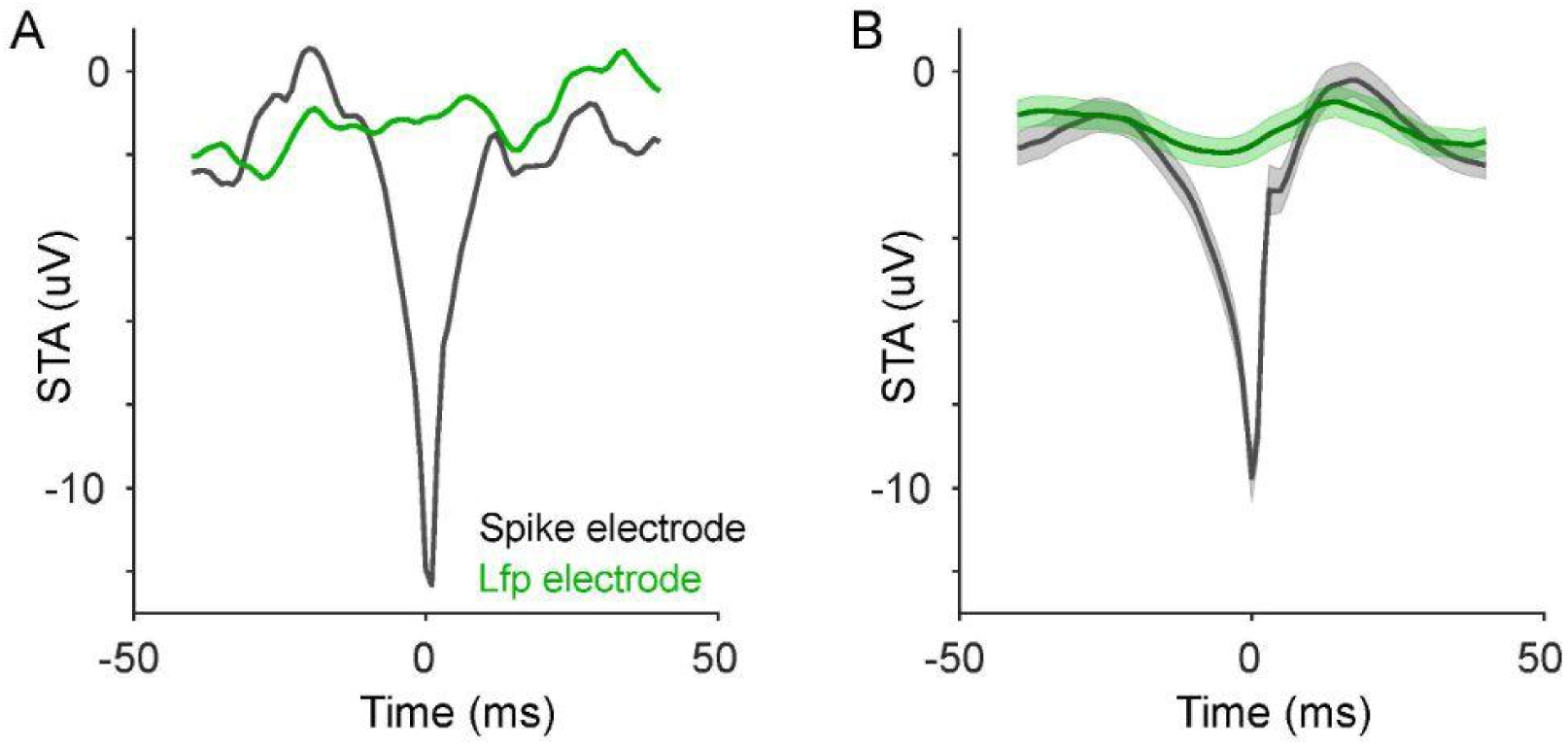
**(A)** Spike triggered average (STA) for an example VM neuron recording. STAs are shown for the spiking electrode and neighboring electrode (∼1.5 mm apart). **(B)** Population average STA of VM neurons (N=139) for spiking and LFP electrodes. The s.e.m. of STA is shown in lighter shades.

## Notes

### Competing Interest Statement

The authors have declared no competing interest.

## References

Anderson, M.C., and Weaver, C. (2009). Inhibitory Control over Action and Memory. Encyclopedia of Neuroscience 153–163. https://doi.org/10.1016/b978-008045046-9.00421-6.

Anderson, B.A., Laurent, P.A., and Yantis, S. (2011). Value-driven attentional capture. Proc. Natl. Acad. Sci. U. S. A. 108, 10367–10371. .

Antzoulatos, E.G., and Miller, E.K. (2016). Synchronous beta rhythms of frontoparietal networks support only behaviorally relevant representations. Elife 5. https://doi.org/10.7554/eLife.17822.

Ardid, S., Vinck, M., Kaping, D., Marquez, S., Everling, S., and Womelsdorf, T. (2015). Mapping of functionally characterized cell classes onto canonical circuit operations in primate prefrontal cortex. J. Neurosci. 35, 2975–2991. .

Awh, E., Armstrong, K.M., and Moore, T. (2006). Visual and oculomotor selection: links, causes and implications for spatial attention. Trends Cogn. Sci. 10, 124–130. .

Bastos, A.M., Vezoli, J., Bosman, C.A., Schoffelen, J.-M., Oostenveld, R., Dowdall, J.R., De Weerd, P., Kennedy, H., and Fries, P. (2015). Visual areas exert feedforward and feedback influences through distinct frequency channels. Neuron 85, 390–401. .

Benjamini, Y., and Hochberg, Y. (1995). Controlling the false discovery rate: A practical and powerful approach to multiple testing. J. R. Stat. Soc. 57, 289–300. .

Bisley, J.W., and Goldberg, M.E. (2003). Neuronal activity in the lateral intraparietal area and spatial attention. Science 299, 81–86. .

Buckley, M.J., Mansouri, F.A., Hoda, H., Mahboubi, M., Browning, P.G.F., Kwok, S.C., Phillips, A., and Tanaka, K. (2009). Dissociable components of rule-guided behavior depend on distinct medial and prefrontal regions. Science 325, 52–58. .

Buschman, T.J., and Kastner, S. (2015). From Behavior to Neural Dynamics: An Integrated Theory of Attention. Neuron 88, 127–144. .

Buschman, T.J., and Miller, E.K. (2007). Top-down versus bottom-up control of attention in the prefrontal and posterior parietal cortices. Science 315, 1860–1862. .

Buschman, T.J., Denovellis, E.L., Diogo, C., Bullock, D., and Miller, E.K. (2012). Synchronous oscillatory neural ensembles for rules in the prefrontal cortex. Neuron 76, 838–846. .

Carrasco, M. (2011). Visual attention: the past 25 years. Vision Res. 51, 1484–1525. .

Carrasco, M., Ling, S., and Read, S. (2004). Attention alters appearance. Nat. Neurosci. 7, 308–313. .

Cassim, F., Monaca, C., Szurhaj, W., Bourriez, J.L., Defebvre, L., Derambure, P., and Guieu, J.D. (2001). Does post-movement beta synchronization reflect an idling motor cortex? Neuroreport 12, 3859–3863. .

Chen, X., Zirnsak, M., Vega, G.M., Govil, E., Lomber, S.G., and Moore, T. (2020). Parietal Cortex Regulates Visual Salience and Salience-Driven Behavior. Neuron 106, 177–187.e4. .

Corbetta, M., and Shulman, G.L. (2002). Control of goal-directed and stimulus-driven attention in the brain. Nat. Rev. Neurosci. 3, 201–215. .

Corbetta, M., Akbudak, E., Conturo, T.E., Snyder, A.Z., Ollinger, J.M., Drury, H.A., Linenweber, M.R., Petersen, S.E., Raichle, M.E., Van Essen, D.C., et al. (1998). A common network of functional areas for attention and eye movements. Neuron 21, 761–773. .

Dean, H.L., Hagan, M.A., and Pesaran, B. (2012). Only coherent spiking in posterior parietal cortex coordinates looking and reaching. Neuron 73, 829–841. .

Dugué, L., Merriam, E.P., Heeger, D.J., and Carrasco, M. (2020). Differential impact of endogenous and exogenous attention on activity in human visual cortex. Sci. Rep. 10, 21274. .

Fiebelkorn, I.C., and Kastner, S. (2019). A Rhythmic Theory of Attention. Trends Cogn. Sci. 23, 87–101. .

Fiebelkorn, I.C., and Kastner, S. (2020). Functional Specialization in the Attention Network. Annu. Rev. Psychol. 71, 221–249. .

Fiebelkorn, I.C., and Kastner, S. (2021). Spike Timing in the Attention Network Predicts Behavioral Outcome Prior to Target Selection. Neuron 109, 177–188.e4. .

Fiebelkorn, I.C., Pinsk, M.A., and Kastner, S. (2018). A Dynamic Interplay within the Frontoparietal Network Underlies Rhythmic Spatial Attention. Neuron 99, 842–853.e8. .

Funahashi, S., Bruce, C.J., and Goldman-Rakic, P.S. (1989). Mnemonic coding of visual space in the monkey’s dorsolateral prefrontal cortex. J. Neurophysiol. 61, 331–349. .

Gottlieb, J.P., Kusunoki, M., and Goldberg, M.E. (1998). The representation of visual salience in monkey parietal cortex. Nature 391, 481–484. .

Gregoriou, G.G., Rossi, A.F., Ungerleider, L.G., and Desimone, R. (2014). Lesions of prefrontal cortex reduce attentional modulation of neuronal responses and synchrony in V4. Nat. Neurosci. 17, 1003–1011. .

Hagan, M.A., and Pesaran, B. (2022). Modulation of inhibitory communication coordinates looking and reaching. Nature 604, 708–713. .

Hagan, M.A., Dean, H.L., and Pesaran, B. (2012). Spike-field activity in parietal area LIP during coordinated reach and saccade movements. J. Neurophysiol. 107, 1275–1290. .

Hanslmayr, S., Matuschek, J., and Fellner, M.-C. (2014). Entrainment of prefrontal beta oscillations induces an endogenous echo and impairs memory formation. Curr. Biol. 24, 904–909. .

Jahfari, S., and Theeuwes, J. (2017). Sensitivity to value-driven attention is predicted by how we learn from value. Psychon. Bull. Rev. 24, 408–415. .

Kaping, D., Vinck, M., Hutchison, R.M., Everling, S., and Womelsdorf, T. (2011). Specific contributions of ventromedial, anterior cingulate, and lateral prefrontal cortex for attentional selection and stimulus valuation. PLoS Biol. 9, e1001224. .

Kilavik, B.E., Ponce-Alvarez, A., Trachel, R., Confais, J., Takerkart, S., and Riehle, A. (2012). Context-related frequency modulations of macaque motor cortical LFP beta oscillations. Cereb. Cortex 22, 2148–2159. .

Kilavik, B.E., Zaepffel, M., Brovelli, A., MacKay, W.A., and Riehle, A. (2013). The ups and downs of β oscillations in sensorimotor cortex. Exp. Neurol. 245, 15–26. .

Knudsen, E.B., and Wallis, J.D. (2020). Closed-Loop Theta Stimulation in the Orbitofrontal Cortex Prevents Reward-Based Learning. Neuron 106, 537–547.e4. .

Kowler, E., Anderson, E., Dosher, B., and Blaser, E. (1995). The role of attention in the programming of saccades. Vision Res. 35, 1897–1916. .

Leon, M.I., and Shadlen, M.N. (1999). Effect of expected reward magnitude on the response of neurons in the dorsolateral prefrontal cortex of the macaque. Neuron 24, 415–425. .

Lisman, J.E. (1997). Bursts as a unit of neural information: making unreliable synapses reliable. Trends Neurosci. 20, 38–43. .

Liu, T., Abrams, J., and Carrasco, M. (2009). Voluntary attention enhances contrast appearance. Psychol. Sci. 20, 354–362. .

Lundqvist, M., Herman, P., Warden, M.R., Brincat, S.L., and Miller, E.K. (2018). Gamma and beta bursts during working memory readout suggest roles in its volitional control. Nat. Commun. 9, 394. .

Maris, E., Schoffelen, J.-M., and Fries, P. (2007). Nonparametric statistical testing of coherence differences. J. Neurosci. Methods 163, 161–175. .

Markowitz, D.A., Shewcraft, R.A., Wong, Y.T., and Pesaran, B. (2011). Competition for visual selection in the oculomotor system. J. Neurosci. 31, 9298–9306. .

Markowitz, D.A., Curtis, C.E., and Pesaran, B. (2015). Multiple component networks support working memory in prefrontal cortex. Proc. Natl. Acad. Sci. U. S. A. 112, 11084–11089. .

Miller, E.K. (2000). The prefrontal cortex and cognitive control. Nat. Rev. Neurosci. 1, 59–65.

Miller, E.K., and Cohen, J.D. (2001). An integrative theory of prefrontal cortex function. Annu. Rev. Neurosci. 24, 167–202. .

Miller, E.K., Lundqvist, M., and Bastos, A.M. (2018). Working Memory 2.0. Neuron 100, 463–475. .

Mitra, P.P., and Pesaran, B. (1999). Analysis of dynamic brain imaging data. Biophys. J. 76, 691–708. .

Moore, T., and Armstrong, K.M. (2003). Selective gating of visual signals by microstimulation of frontal cortex. Nature 421, 370–373. .

Moore, T., and Zirnsak, M. (2017). Neural Mechanisms of Selective Visual Attention. Annu. Rev. Psychol. 68, 47–72. .

Paneri, S., and Gregoriou, G.G. (2017). Top-Down Control of Visual Attention by the Prefrontal Cortex. Functional Specialization and Long-Range Interactions. Front. Neurosci. 11, 545. .

Panichello, M.F., and Buschman, T.J. (2021). Shared mechanisms underlie the control of working memory and attention. Nature 592, 601–605. .

Pape, A.-A., and Siegel, M. (2016). Motor cortex activity predicts response alternation during 40 sensorimotor decisions. Nat. Commun. 7, 13098. .

Pesaran, B., Pezaris, J.S., Sahani, M., Mitra, P.P., and Andersen, R.A. (2002). Temporal structure in neuronal activity during working memory in macaque parietal cortex. Nat. Neurosci. 5, 805–811. .

Pesaran, B., Vinck, M., Einevoll, G.T., Sirota, A., Fries, P., Siegel, M., Truccolo, W., Schroeder, C.E., and Srinivasan, R. (2018). Investigating large-scale brain dynamics using field potential recordings: analysis and interpretation. Nat. Neurosci. 21, 903–919. .

Pesaran, B., Hagan, M., Qiao, S., and Shewcraft, R. (2021). Multiregional communication and the channel modulation hypothesis. Curr. Opin. Neurobiol. 66, 250–257. .

Petrides, M. (2005). Lateral prefrontal cortex: architectonic and functional organization. Philos. Trans. R. Soc. Lond. B Biol. Sci. 360, 781–795. .

Petrides, M., and Pandya, D.N. (1994). Comparative architectonic analysis of the human and the macaque frontal cortex. Handbook of Neuropsychology 17–58. .

Phillips, J.M., Vinck, M., Everling, S., and Womelsdorf, T. (2014). A long-range fronto-parietal 5- to 10-Hz network predicts “top-down” controlled guidance in a task-switch paradigm. Cereb. Cortex 24, 1996–2008. .

Reynolds, J.H., Chelazzi, L., and Desimone, R. (1999). Competitive mechanisms subserve attention in macaque areas V2 and V4. J. Neurosci. 19, 1736–1753. .

Rossi, A.F., Bichot, N.P., Desimone, R., and Ungerleider, L.G. (2007). Top down attentional deficits in macaques with lesions of lateral prefrontal cortex. J. Neurosci. 27, 11306–11314. .

Rushworth, M.F.S., Buckley, M.J., Gough, P.M., Alexander, I.H., Kyriazis, D., McDonald, K.R., and Passingham, R.E. (2005). Attentional selection and action selection in the ventral and orbital prefrontal cortex. J. Neurosci. 25, 11628–11636. .

Salazar, R.F., Dotson, N.M., Bressler, S.L., and Gray, C.M. (2012). Content-specific fronto-parietal synchronization during visual working memory. Science 338, 1097–1100. .

Sanes, J.N., and Donoghue, J.P. (1993). Oscillations in local field potentials of the primate motor cortex during voluntary movement. Proc. Natl. Acad. Sci. U. S. A. 90, 4470–4474. .

Schmidt, R., Herrojo Ruiz, M., Kilavik, B.E., Lundqvist, M., Starr, P.A., and Aron, A.R. (2019). Beta Oscillations in Working Memory, Executive Control of Movement and Thought, and Sensorimotor Function. J. Neurosci. 39, 8231–8238. .

Spitzer, B., and Haegens, S. (2017). Beyond the Status Quo: A Role for Beta Oscillations in Endogenous Content (Re)Activation. eNeuro 4. https://doi.org/10.1523/ENEURO.0170-17.2017.

Staudigl, T., Minxha, J., Mamelak, A.N., Gothard, K.M., and Rutishauser, U. (2022). Saccade-related neural communication in the human medial temporal lobe is modulated by the social relevance of stimuli. Sci Adv 8, eabl6037. .

Suzuki, M., and Gottlieb, J. (2013). Distinct neural mechanisms of distractor suppression in the frontal and parietal lobe. Nat. Neurosci. 16, 98–104. .

Theeuwes, J. (2010). Top-down and bottom-up control of visual selection. Acta Psychol. 135, 77–99. .

Voloh, B., and Womelsdorf, T. (2016). A Role of Phase-Resetting in Coordinating Large Scale Neural Networks During Attention and Goal-Directed Behavior. Front. Syst. Neurosci. 10, 18. .

Womelsdorf, T., and Everling, S. (2015). Long-Range Attention Networks: Circuit Motifs Underlying Endogenously Controlled Stimulus Selection. Trends Neurosci. 38, 682–700. .

Womelsdorf, T., Ardid, S., Everling, S., and Valiante, T.A. (2014). Burst firing synchronizes prefrontal and anterior cingulate cortex during attentional control. Curr. Biol. 24, 2613–2621.

